# Lysine Potentiates Insulin Secretion via AASS-Dependent Catabolism and Regulation of GABA Content and Signaling

**DOI:** 10.1101/2025.03.03.641187

**Authors:** Felipe Munoz, Qian Gao, Matthias Mattanovich, Kajetan Trost, Ondřej Hodek, Andreas Lindqvist, Nils Wierup, Malin Fex, Thomas Moritz, Hindrik Mulder, Luis Rodrigo Cataldo

## Abstract

**Background and aim:** Lysine is an essential amino acid with insulinotropic effects in humans. *In vitro*, lysine also potentiates glucose-stimulated insulin secretion (GSIS) in β cell lines and rodent pancreatic islets. For decades it has been assumed that insulinotropic action of lysine is mediated by plasma membrane depolarization similar to arginine. Aminoadipate-Semialdehyde Synthase (AASS) is a mitochondrial-located bifunctional enzyme engaged in the first two steps of the lysine catabolism. Whether AASS-dependent lysine catabolism occurs in β cells and whether it is required for its insulinotropic action has not been investigated.

**Methods:** mRNA expression of lysine catabolism pathway genes was assessed in human islets from non-diabetic (ND) and type 2 diabetes (T2D) subjects. AASS was silenced in human pancreatic islets and in INS1 832/13 β cells. β cell metabolism and function were investigated by ELISA, extracellular flux analysis, live cell calcium imaging, transcriptomics and metabolomics analyses.

**Results:** Expression of genes involved in lysine catabolism, including *AASS, ALDH7A1, DHTKD1* and *HADH*, was reduced in pancreatic islets from T2D donors. Silencing of *AASS* resulted in reduced lysine- and glucose-stimulated insulin secretion in human islets and INS1 832/13 β cells. Surprisingly, transcriptomics and metabolomics analysis in *Aass*-KD β cells with suppressed lysine catabolism identified reduced γ-aminobutyric acid (GABA)/glutamate ratio as well as altered expression of genes implicated in GABA metabolism. This was accompanied by altered mitochondrial TCA cycle and oxidative phosphorylation (OXPHOS) activity, reflected by elevated lactate/pyruvate and reduced whole-cell ATP/ADP content as well as ATP-linked mitochondrial respiration. Glucose-and GABA-stimulated cytosolic calcium was also altered in *Aass-KD* β cells. Strikingly, addition of GABA recovered impaired insulin secretion in *Aass-KD* β cells.

**Conclusion:** AASS-dependent lysine catabolism is required to maintain adequate GABA shunt metabolism and signaling. In addition, lysine catabolism supports mitochondrial energy production, calcium uptake and insulin secretion. Reduced AASS-dependent lysine catabolism may contribute to β cell GABA depletion and dysfunction in T2D patients.

## 1. Introduction

A recent systematic analysis of insulin secretion dynamics in human pancreatic islets in response to macronutrients, highlighted the insulinotropic actions of amino acids^1^. Notably, insulin secretion in islets from 9% of donors was more strongly stimulated by amino acids than by glucose^1^. This study highlights the necessity of further investigating the mechanisms by which non-glucose nutrients, including amino acids, influence metabolic responses in human islets and their role in regulating glucose homeostasis.

Lysine is an essential amino acid with established insulinotropic effects in humans for over 50 years^2^. A study administering equal intravenous doses (30 g) of individual amino acids in healthy subjects found that lysine has greater insulinotropic potency than leucine and comparable to arginine^2^. In line with this, the strong insulinogenic index of some dairy food with low glycaemic index has been attributed to raised plasma levels of lysine, arginine and branched chain amino acids^3^.

The ability of lysine to directly potentiate insulin secretion in β cells has also been demonstrated by *in vitro* studies. The potency of amino acids to enhance insulin secretion in the presence of glucose was compared in INS1E cells and mouse islets by administrating equimolar concentrations (10 mM) of each amino acid. Lysine and arginine were the most potent amino acids to enhance GSIS^4^.

It is well-established that several amino acids potentiate insulin secretion in the presence of glucose^5^. The mechanisms of action of some amino acids are metabolism-dependent, where they act as energy substrates and increase the production of ATP. The mechanism of other amino acids is metabolism-independent and insulin secretion is potentiated by an electrochemical effect, after depolarizing the plasma membrane via their positively charged properties or upon co-transporting Na^+^^5,6^. Due to its positive charge at physiological pH, similar to arginine, it’s been assumed for decades that the insulinotropic action of lysine is mediated by its uptake in β cells, leading to plasma membrane depolarization^4^. However, it remains unclear whether the insulinotropic effect of lysine is mediated by an electrochemical action or by a metabolism-dependent mechanism. Furthermore, the potential contribution of lysine metabolism within β cells to the amplification of GSIS has not been investigated.

Aminoadipate-Semialdehyde Synthase (AASS) is a mitochondrially-located bifunctional enzyme with lysine-ketoglutarate reductase (LKR) and saccharopine dehydrogenase (SDH) activities^7,8^, carrying out the first two steps in the saccharopine catabolic pathway of lysine. AASS-mediated lysine catabolism may directly potentiate insulin secretion since insulinotropic metabolites such as glutamate and acetyl-CoA are produced along the pathway. Indeed, GC-MS-based labelling experiments have shown that lysine is the precursor for *de novo* synthesis of up to 33% of the glutamate pool in the mammalian brain^7^. Glutamate is accumulated in β cells in response to glucose stimulation and enhances GSIS, hence considered a Metabolic Coupling Factor (MCF)^9,10^. Glutamate anaplerotically enters the TCA cycle upon conversion to α-ketoglutarate by the enzyme glutamate dehydrogenase^9^ or exits mitochondria and is taken up into insulin granules to promote exocytosis^9,11^. In addition, glutamate may represent a route for γ-aminobutyric acid (GABA) provision as glutamate is a direct precursor for GABA synthesis in β cells^12^, another important regulator of β cell mass and function^13, 14^. On the other hand, acetyl-CoA, an end product of the AASS-dependent lysine catabolism can increase GSIS by feeding carbons into the TCA cycle or be exported from mitochondria to the cytosol as a substrate for acetyl-CoA carboxylase (ACC) to produce malonyl-CoA in the cytosol, another MCF for GSIS^10^.

The lysine catabolism pathway holds significant clinical relevance, as epidemiological studies have identified plasma levels of lysine and its metabolites, such as 2-aminoadipic acid, as predictive risk factors for type 2 diabetes (T2D)^15,16^. Furthermore, mutations in genes encoding enzymes involved in this pathway, including *AASS*, have been implicated in inborn errors of metabolism, such as hyperlysinemia^17^. Most of these metabolic disorders are associated with accumulation of toxic levels of metabolites produced along the lysine catabolic pathway. Therefore, AASS-dependent catabolism of lysine may play a dual role in β cell function by maintaining lysine at physiological levels and providing lysine-derived metabolites that amplifies GSIS.

## 2. Research Design and Methods

### 2.1 Human Pancreatic Islet Cohort

Human pancreatic islets were obtained from the EXODIAB Human Tissue Laboratory, which receives islets from the Nordic Network for Clinical Islet Transplantation (http://www.nordicislets.org). The clinical characteristics of the donors of islets from this cohort (n = 188) and all methodological details related to RNA sequencing of islets have been published^18^.

### 2.2 Cell culture

INS1 832/13 and EndoC-βH1 cells were cultured as previously described^19,20^. EndoC-βH1 cells were cultured in Matrigel/fibronectin-coated (100/2 mg/mL, Sigma-Aldrich) flasks with DMEM containing 5.6 mmol/L glucose, 2% BSA, 10 mmol/L nicotinamide, 50 mmol/L β-mercaptoethanol, 5.5 mg/mL transferrin, 6.7 ng/mL sodium selenite, 100 IU/mL penicillin, and 100 mg/mL streptomycin at 37°C in a humidified atmosphere with 5% CO_2_.

### 2.3 GSIS in **β** cell lines and human islets

INS1 832/13 cells (1.7×10^5^ cells/cm^2^) and EndoC-βH1 cells (1.5×10^5^ cells/cm^2^) were seeded in 24 and 48 well-plates, respectively, 96 hours prior to experiments. Seeded cells and human pancreatic islets (10 islets/well, 24-WP, 4-6 replicates/condition) were starved in secretion assay buffer (SAB) buffer (final SAB 1X composition was (in mM): 114 NaCl; 4.7 KCl; 1.2 KH2PO4; 1.16 MgSO4; 20 HEPES; 2.5 CaCl2; 25.5 NaHCO3; 0.2% bovine serum albumin (BSA), pH 7.2-7.4) with low glucose (LG, 1-2.8 mM mM). Then, cells and islets were simultaneously (in parallel wells) stimulated for 1h with SAB LG, high glucose (HG, 16.7-20mM) or HG+3-isobutyl-1-methylxanthine (IBMX) (100µM). The insulin concentration in the supernatants (accumulated during 1h of stimulation) was measured using the insulin ELISA: for samples from INS1 832/13 cells (non-diluted) High Range Rat Insulin ELISA (Mercodia, catalogue number (#): 10-1145-01), for samples from EndoC-βH1 cells (dilution factor: 12.5) and from human islets (dilution factor: 5) the Human Insulin Elisa (Mercodia, #10-1113-10). The total insulin content was measured in EndoC-βH1 cells and human islets by extraction of total cells/islet protein with RIPA buffer (composition was: 150 mM NaCl, 1% NP40, 0.5% sodium deoxycholate, 0.1% SDS, 50 mM TRIS-HCl, pH 8.0); insulin secretion was normalized to the total insulin content and expressed as % secreted insulin.

### 2.4 AASS silencing in **β** cell lines and human islets

INS1 832/13 cells (1.5×10^5^ cells/cm^2^) were seeded in 24 well plates and transfected after 24 h with 10 nM of scramble negative control (NC) (Thermofisher, custom select siRNA: 5’-GAGACCCUAUCCGUGAUUAUU-3’) or rat *Aass* siRNA (Thermofisher, #4390771; s150503). EndoC-βH1 cells (1.8×10^5^ cells/cm^2^) were seeded in 48 well plates and transfected 24 and 48 h after (two successive days, double shot) with 40 nM of Silencer Select Negative Control No. 1 siRNA (Thermofisher, #4390843) or human *AASS* siRNA (Thermofisher, # 4392420; s19785). Human pancreatic islets (400-500) were seeded in 35 mm petri dish containing 2 ml of RPMI medium (5mM glucose, 10% FBS (Sigma #7524), 200 mM L-Glutamine) and transfected on two successive days with 0.5 ml of 50 nM of a Silencer Select Negative Control No. 1 siRNA (Thermofisher, #4390843) or human *AASS* siRNA (Thermofisher, #4392420; s19785), using Lipofectamine RNAiMAX Transfection Reagent (Thermofisher, #13778075). Details of donors of pancreatic islets used for experiments of *AASS* silencing are given in Table 1 (Fig. S1A). Human pancreatic islets were cultured as previously described^21^. All procedures were approved by the Swedish Ethical Review Authority.

### 2.5 mRNA expression analysis

Total RNA was isolated by RNeasy Mini Kit (Qiagen) and cDNA was generated with RevertAid First-Strand cDNA synthesis kit. Ten ng cDNA/well (in duplicates) were used for qPCR; *Aass* rat (Rn01455736_m1) and human *AASS* (Hs00194991_m1) taqman assays were obtained from Thermofisher. qPCR assays were performed using Applied Biosystems QuantStudio7 Flex Real-Time PCR System (Thermofisher). Relative gene abundance was calculated using the ΔΔCt method with β*-actin* as reference gene and expressed as fold change to control.

### 2.6 Lactate measurement

INS1 832/13 cells (1.7×10^5^ cells/cm^2^) were seeded in 24 well plates, transfected with scramble (negative control, NC) or *Aass* siRNA (10 nM) (as described in section 2.3) and GSIS determined (as described in section 2.2). The concentration of the extracellular lactate accumulated during 1h in response to stimulation with SAB buffer (LG or HG) was measured by use of a Lactate Colorimetric/Fluorometric Assay Kit (Biovision, #607-100) following the manufacturer’s instructions. Briefly, cell conditioned samples and lactate standards were mixed with an enzyme plus fluorescent probe mix in a ratio 1:1, incubated for 30 min (at room temperature) and fluorescence emission read (Ex/Em = 535/590 nm) in a microplate reader. Lactate concentrations were normalized by total cell mass in respective wells (protein mass measured by Pierce BCA Protein Assay Kit).

### 2.7 Metabolomics analysis

The cellular metabolome was analyzed by combined gas chromatography-mass spectrometry (GC-MS) as previously described^22,23^. Briefly, INS1 832/13 cells were stimulated with SAB LG, HG or HG+IBMX for 1h (as described for GSIS above) and total cellular metabolites extracted with methanol/water (9/1, v/v) containing a cocktail of stable isotope labelled internal standards (^2^H_4_-succinate (0.22 mg/L), ^2^H_8_-valine (4.41 mg/L), ^13^C_4_-3-hydroxybutyric acid (2.21 mg/L), ^13^C_5_^15^N-glutamic acid (22.06 mg/L), ^2^H_19_-decanoic acid (4.41 mg/L) and ^2^H_31_-palmitic acid (22.06 mg/L). Proteins were precipitated by three repeated rounds of shaking/snap freezing on liquid nitrogen, incubation in ice for 1 h and centrifugation. Seven independent experiments were performed. The generated raw data were pre-processed using an in-house script developed at Swedish Metabolomics Centre, Umeå, Sweden. The detected peaks were identified by comparison of mass spectra and retention indexes using NIST MS Search v.2.0, using in-house and NIST98 spectral databases. After pre-processing and filtering of the metabolites of high coefficient of variation in quality control pool samples (>25%), levels of 21 metabolites were used for further analysis.

### 2.8 Mitochondrial oxygen consumption (OCR) measurements

OCR was evaluated with a Seahorse XFe24 extracellular flux analyzer (Agilent Technologies). INS1 832/13 cells were transfected with scramble (negative control, NC) or *Aass* siRNA (10 nM) and cultured for 72h. Cells were starved in SAB LG (2.8 mM) for 2h and OCR was measured in SAB (without bicarbonate/HEPES) every 3 min for 90 min. OCR was measured at basal glucose (2.8mM glucose) and after adding HG (16.7mM), 5μM oligomycin, 4μM Carbonyl cyanide p-trifluoromethoxyphenylhydrazone (FCCP), and 1μM rotenone/antimycin A. Wave Seahorse Software and an online tool (seahorseanalytics.agilent.com) were used to analyze data. Non-mitochondrial respiration was subtracted, and OCR data normalized to total protein content, determined by the colorimetric Biuret method (Pierce-BCA Protein Assay Kit, Thermofisher).

### 2.9 Quantification of whole-cell ATP and ADP

Cellular ATP and ADP levels were quantified by ultra-high performance liquid chromatography-tandem mass spectrometry (UHPLC-MS/MS). Prior to analysis, the cell extracts were diluted 10 times with MeOH/water (1/1, v/v) with final concentration of the labelled internal standards (AMP-^13^C_10_^15^N_5_, ADP-^15^N_5_, ATP-^13^C_10_) of 1 µM. Separation of the nucleotides was achieved with a 15-minute gradient using a iHILIC-(P) Classic column (PEEK, 50 × 2.1 mm, 5 µm, HILICON, Umeå, Sweden) with mobile phases composed of (A) 10 mM ammonium acetate in water at pH 9.4 and (B) 10 mM ammonium acetate at pH 9.4 in 90% acetonitrile, both mobile phases were supplemented with 5 µM medronic acid. The flow rate was 0.35 mL/min, and the gradient elution program was set as follows: 0.0 min (85% B), 5 min (60% B), 7 min (30% B), 8 min (30% B), 9 min (85% B), 15 min (85% B). The UHPLC-MS/MS system consisted of an Agilent 1290 UPLC connected to an Agilent 6490 triple quadrupole tandem mass spectrometer (Agilent, CA, USA). Analytes were ionized using electrospray ionization operated in positive ionization mode. The source and gas parameters were set as follows: ion spray voltage 4.0 kV, gas temperature 150°C, drying gas flow 11 L/min, nebulizer pressure 20 psi, sheath gas temperature 325°C, sheath gas flow rate 12 L/min, fragmentor 380 V. Multiple reaction monitoring (MRM) transitions for AMP, ADP, ATP, and their respective labelled internal standards were optimized by flow injection analysis. Quantification of AMP, ADP, and ATP was conducted based on internal standard calibration. Calibration curves were linear from 5 nM to 50 µM. The accuracy, determined through spiking experiments of the cell extracts (n = 4) with 10 µM standards, was within the acceptable range of 100 ± 15%.

### 2.10 Quantification of extracellular GABA and glutamate

Cellular GABA and glutamate levels were quantified by ultra-high performance liquid chromatography-tandem mass spectrometry (UHPLC-MS/MS). 100 µl of cellular extracts including 100 ng of nor-valine as internal standard was evaporated to dryness. Extracted samples were derivatized by AccQ-Tag™ (Waters, Milford, MA, USA) according to the manufacturer’s instructions. Briefly, the dry extract was dissolved in 20 µl 20mM HCl, 60 μL of AccQ•Tag Ultra Borate buffer and 20 μL of the freshly prepared AccQ•Tag derivatization solution and the sample was immediately vortex for 30s.

Samples were kept at room temperature for 30 minutes followed by 10 minutes at 55°C. The quantification of the amino acids was achieved on the liquid chromatography-tandem mass spectrometer consisting of a Waters Acquity UPLC I-Class connected to a Waters Xevo TQ-XS tandem mass spectrometer (Waters, Manchester, UK). The separation was performed by injecting 1 µL of each sample to an Acquity UPLC BEH C18 column (100×2.1 mm, 1.7 µm, Waters, MA, USA). The mobile phase consisted of (A) 0.1% formic acid and (B) acetonitrile with 0.1% formic acid and was delivered on the column by a flow rate of 0.50 mL/min with the following gradient: The initial conditions consisted of 0.1% B, and the following gradient was used with linear increments: 0.54-3.50 minutes (0.1-9.1% B), 3.50-7.0 (9.1-17.0% B), 7.0-8.0 (17.0-19.70% B), 8.0-8.5 (19.7% B), 8.5-9.0 (19.7-21.2% B), 9.0-10.0 (21.2-59.6% B), 10.0-11.0 (59.6-95.0% B), 11.0-11.5 (95.0% B), 11.5-15.0 (0.1% B). Column and autosampler were thermostated at 55 °C and 4 °C, respectively. Analytes were ionized in an electrospray ion source operated in the negative mode. The source and gas parameters were set as follows: ion spray voltage 3.5 kV, desolvation temperature 300 °C, desolvation gas flow 800 L/hr, nebulizer pressure 7 Bar, cone gas flow 150 L/h. The instrument was operated in multiple reaction monitoring mode (MRM), and dwell time was set to 30 ms for all transitions. For quantification 8-points calibration curves were used including different levels of non-labelled and constant levels of the labelled nor-valine internal standard. The instrument was controlled by MassLynx 4.2, and data processing was performed with TargetLynx XS (Waters, MA, USA).

### 2.11 mRNA sequencing analysis

Messenger RNA sequencing (mRNAseq) was performed by the Single-Cell Omics platform (SCOP) at the Novo Nordisk Foundation Centre for Basic Metabolic Research. Total RNA was isolated using the RNeasy Mini Kit (Qiagen) according to manufacturer’s protocol from NC and *Aass*-KD INS1 832/13 cells. Total RNA was treated with DNAse and delivered to the SCOP where they performed RNA quality control analysis (20 ng of RNA) and prepared RNA-seq library (250 ng of RNA). Libraries were prepared using the Universal Plus mRNA-seq protocol (Tecan, CH) as recommended by the manufacturer. Libraries were quantified with NuQuant using the CLARIOstar Plate Reader (BMG Labtech, DE), quality checked using a TapeStation instrument (Agilent Technologies, US) and subjected to 52-bp paired-end sequencing on a NovaSeq 6000 (Illumina, US).

For the RNAseq data analysis, the nf-core^24^ RNA-seq pipeline was used to align RNA-seq reads against the rat genome assembly and *Aass* transcripts^25^. Testing for differential expression was performed using edgeR^26^ v.3.38.4, using the quasi-likelihood framework with a fitted model of the form ∼ 0 + group, where group encoded both genotype and treatment. Contrasts were constructed as described in the edgeR manual. Gene Ontology^27^ and Reactome^28^ enrichments were found using the CAMERA function^29^ which is part of the edgeR package. Only gene ontologies with between 5 and 500 genes were investigated.

### 2.12 Statistical analysis

Data are shown as mean±SEM of at least three independent experiments (unless specified). The Student’s t test was used to compare two groups and ANOVA for multiple groups. Statistical significance was set at p<0.05. SPSS 28 and Prism 9.0 (GraphPad) were used for correlational and basic statistical analysis and graph generation. For metabolomics data, batch effects were removed using ComBat^30^ and data expressed as cell content concentration (µM) for each metabolite at LG and HG or as ratio of specified metabolite cell contents. ANOVA was applied for identifying differentiating metabolites between two genotypes and p values were corrected for multiple testing with the Benjamini–Hochberg procedure. Prior to multivariate analysis, data were mean-centered and scaled to unit variance to ensure equal weight of all metabolites in analysis. Unsupervised principal component analysis (PCA) was first used to recognize underlying patterns and to detect outliers. Supervised orthogonal partial least-squares discriminant analysis (OPLS-DA) was then performed to find discriminating metabolites based on the variable importance on projection (VIP) score. The VIP measures the influence of metabolites on the predictive component in the model, in other words; how much influence the metabolite has on discriminating the two different genotypes (*Aass*-KD vs NC control). Metabolites with VIP score > 1 were considered metabolites responsible for the separation between the two genotypes. The OPLS-DA model was validated with 3-fold cross-validation and permutation test (n = 200). All the metabolomics analysis was carried out in R 4.1.2. Pathway enrichment analysis was performed with MetaboAnalyst based on the Kyoto Encyclopedia of Genes and Genomes (KEGG) library using hypergeometric test.

## 3. Results

### 3.1. Effects of lysine on insulin secretion in INS1 832/13 β cells and human pancreatic islets

Intravenous administration of lysine in healthy subjects has insulinotropic effects^2^ and direct stimulation of INS1E cells and mouse islets with lysine enhances GSIS^4^. Therefore, we first aimed to test whether lysine has insulinotropic effects *in vitro* in the glucose-responsive β cell line INS1 832/13 and in human pancreatic islets.

We observed a significant lysine-promoted increase (∼26%) in insulin secretion in the presence of high glucose (HG) (290.7±30.8 vs. 363.6±35.7 ng/mg of protein/h, p-value: 0.0003), but no effects on the presence of low glucose (LG) (Fig. 1A). Similarly, in islets from non-diabetic (ND) donors, acute stimulation with lysine (10 mM) had no effect on insulin secretion at LG, but increased insulin secretion nominally by approximately 25% at HG (6.1±2.5 vs. 7.7±3.1 % of content, p-value: 0.11) (Fig. 1B).

**Figure 1.**
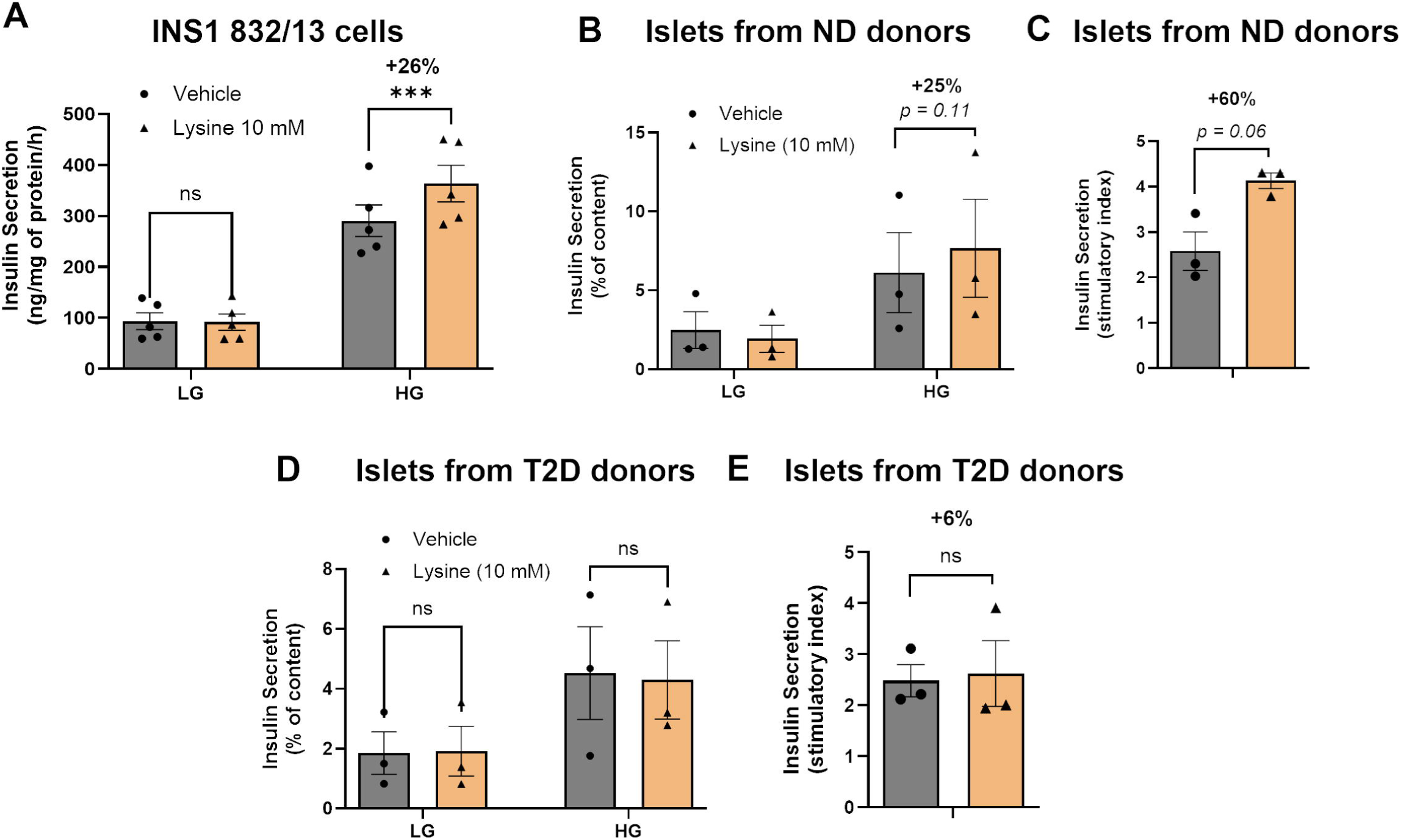
Effects of lysine on insulin secretion in INS1 832/13 β cells and in human pancreatic islets. **A.** Insulin secretion at LG and HG in presence and absence of lysine (10 mM) in INS1 832/13 β cells (n=5). **B, D.** Insulin secretion in human pancreatic islets from ND **(A)** and T2D **(D)** donors at low glucose (LG, 2.8 mM) and high glucose (HG, 16.7 mM) in the absence and presence of lysine (10 mM) (n=3). **C, E.** Calculated lysine-provoked stimulatory index (insulin secretion at HG/LG in presence vs absence of lysine) in ND (**C**) and T2D **(E)** islets (n=3). All panels show individual data points and mean ± s.e.m. Student t-test or Two-way ANOVA (depending on number of compared groups); *p<0.05, **p < 0.01, ***p < 0.001, ****p < 0.0001. ND, non-diabetic; T2D, type 2 diabetes.

We also evaluated the effects of lysine on GSIS by calculating the stimulatory index (the ratio of insulin secretion at HG versus LG) in the presence or absence of lysine. We found a ∼60% nominal increase (p=0.06) in the stimulatory index when islets were stimulated in the presence of lysine (Fig. 1C).

When we conducted the same experiments on pancreatic islets from T2D patients, we observed no significant changes in GSIS in the presence vs. absence of lysine (Fig. 1D, E).

Therefore, these data indicates that lysine potentiates insulin secretion in a glucose dependent manner in INS1 cells and healthy pancreatic islets but not in islets from T2D donors.

### 3.2. Expression of Lysine Catabolism Pathway Genes is Reduced in Pancreatic Islets from T2D Donors, and AASS Expression Levels are Associated with Islet Function

To investigate the relevance of the lysine catabolism pathway in β cell metabolism and insulin secretory function, and whether dysregulation of this pathway contributes to β cell dysfunction during the progression to T2D, we examined the expression of genes encoding the enzymes involved in lysine catabolism (see pathway chart in Fig. 2F) in pancreatic islets from T2D donors (Fig. 2A-E). Interestingly, we found that expression of *AASS* (steps 1 and 2), *ALDH7A1* (step 3), *AADAT* (step 4), *DHTKD1* (step 5), and *HADH* (step 8) was significantly reduced in pancreatic islets from T2D compared to ND donors (Fig. 2A-E). Notably, mRNA expression levels of *ALDH7A1* and *DHTKD1* were highly correlated with *AASS* expression in pancreatic islets (Fig. S1B, C), suggesting a common regulatory mechanism for lysine catabolism pathway genes.

**Figure 2.**
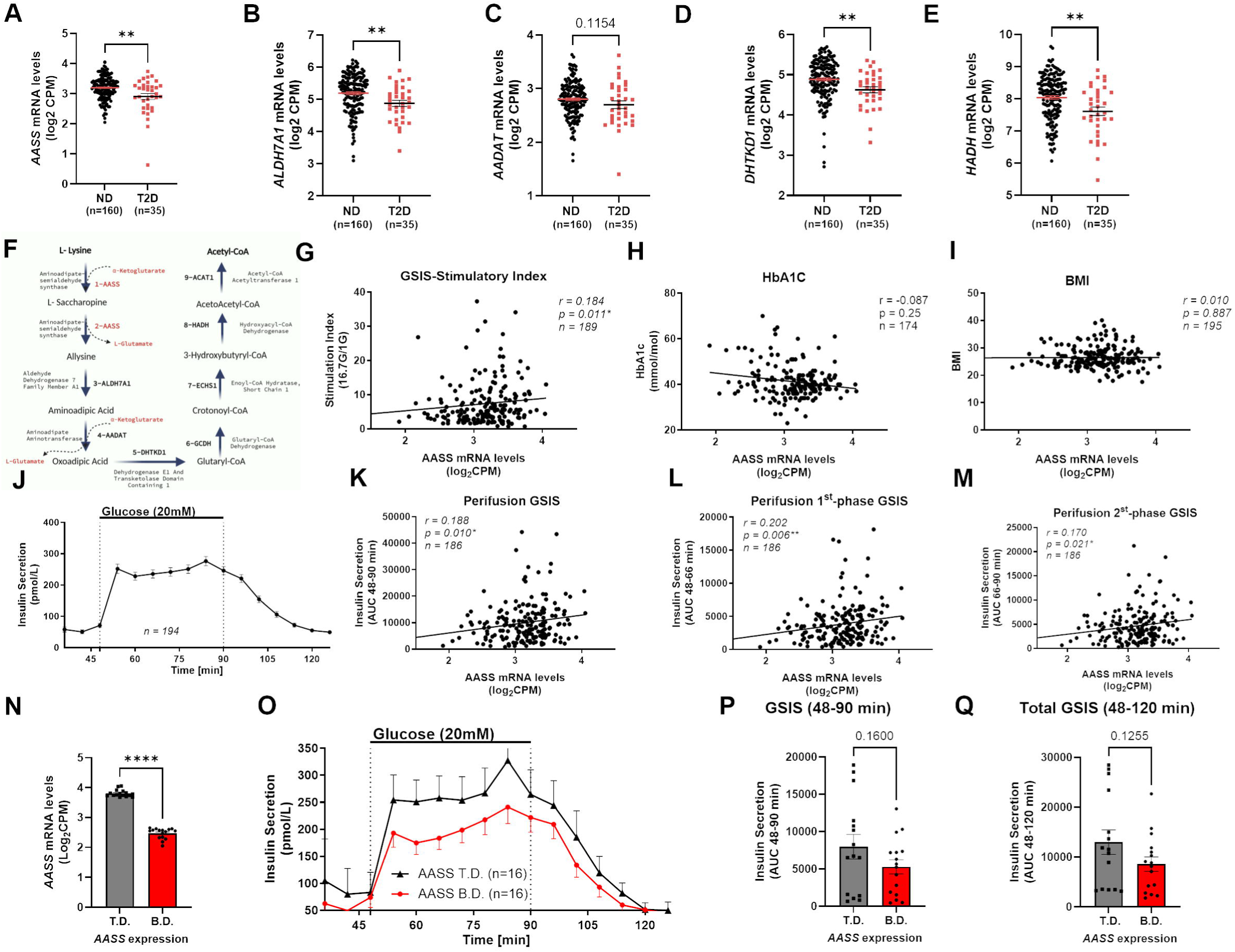
The Expression of Lysine Catabolism Pathway Genes Is Reduced in Pancreatic Islets from T2D Donors, and AASS Expression Levels Are Associated with Islet Function. **A-E.** mRNA levels of lysine catabolism pathway genes in pancreatic islets from ND (n=160) vs. T2D (n=35) donors. **F.** Scheme illustrating the Saccharopine, AASS-dependent lysine catabolism pathway. **G-I.** Spearman rank correlation analyses between *AASS* mRNA levels in islets and **(G)** GSIS stimulatory index (SI), **(H)** donors HbA1c and **I)** donors BMI. **J-M.** Perifusion dynamic GSIS analysis of human islets. **J.** Average dynamic GSIS trace in human pancreatic islets from ND and T2D donors. **K-M.** Spearman rank correlation analyses between *AASS* mRNA levels in islets and **(K)** AUC of total dynamic GSIS, **(L)** AUC of the first-phase GSIS, and **(M)** AUC of the second-phase GSIS. **N.** Average *AASS* mRNA levels in pancreatic islets from ND donors in the lowest decile (B.D.) vs. the highest decile (T.D.). **O.** Traces and **(P, Q)** Averages AUC of dynamic GSIS in islets from ND donors with the B.D. (n=16) vs. the T.D. (n=16) *AASS* mRNA levels. All panels show individual data points and mean ± s.e.m. Student t-test or Two-way ANOVA (depending on number of compared groups); *p<0.05, **p < 0.01, ***p < 0.001, ****p < 0.0001. AASS, Aminoadipate-Semialdehyde Synthase; AUC, area under the curve; GSIS, glucose-stimulated insulin secretion; ND, non-diabetic; T2D, type 2 diabetes.

We also assessed AASS protein expression in pancreatic islets from T2D and ND donors, using immunostaining (Fig. S1D). We confirm the presence of AASS protein in human pancreatic β cells as AASS and INSULIN staining colocalized (Fig. S1D). This is consistent with published transcriptomics data from FACS-sorted human pancreatic islet cells, showing significantly higher AASS mRNA expression in β cells compared to α cells (310±56 vs. 147±39 (normalized counts), q-value: 6e-15)^31^. This finding also aligns with a strong correlation between AASS protein expression and β cell proportion in human islets (Fig. S1E, data obtained from www.humanislets.com). We have not done quantitative AASS protein expression analysis between T2D and ND islets via immunostaining (Fig. S1D), but mass-spectrometry-based proteomics analysis confirmed a significant reduction of AASS protein abundance in T2D compared to ND islets (Fig. S1F, www.humanislets.com).

Since AASS encodes the enzyme responsible for the first two rate-limiting steps in the lysine catabolism pathway^17^, we focused on AASS expression as the key regulator of lysine catabolism. To gain insight into the potential role of AASS-dependent lysine catabolism in human islet function and glucose homeostasis, we evaluated the association between AASS mRNA expression in islets and their *ex vivo* GSIS, glycemic control (HbA1c), and donor BMI. We found that AASS mRNA levels were positively correlated with GSIS (the stimulatory index, SI) (Fig. 2G), but did not correlate with HbA1c (Fig. 2H) or with BMI (Fig. 2I). Importantly, in another cohort of human islets (www.humanislets.com), we confirmed a positive correlation between AASS protein abundance in human islets and GSIS (in response to both 6- or 15-mM glucose) (Fig. S1G, H) and a negative correlation between AASS protein abundance and donor HbA1c (Fig. S1I).

To further explore the former association, we evaluated whether AASS mRNA levels were linked to dynamic GSIS function in islets from ND donors (Fig. 2J). We found a positive correlation between AASS mRNA levels and total insulin secretion (Fig. 2K), as well as with insulin secretion during the 1^st^ phase (Fig. 2L) and 2^nd^ phase (Fig. 2M) of dynamic GSIS.

To further explore the relationship between AASS expression and insulin secretory function, we compared the dynamic GSIS function of islets from ND donors with the lowest (bottom decile (B.D.) (n=16)) and highest (top decile (T.D.) (n=16)) AASS mRNA expression levels (Fig. 2N). We observed that pancreatic islets with the B.D. of AASS expression exhibited reduced insulin secretion during a GSIS perifusion assay (90 min) compared to islets with the T.D. AASS expression (Fig. 2O). The average AUC of insulin secretion during the glucose stimulation period (48-90 min) (Fig. 2P) and throughout the entire assay (48-120 min) (Fig. 2Q) was nominally lower in islets with the B.D. compared to T.D. AASS expression.

Collectively, these findings suggest that pancreatic islets from healthy donors with impaired lysine catabolism, as consequence of reduced *AASS* expression, may lead to reduced insulin secretion. Likewise, impaired lysine catabolism associated to the reduced *AASS* mRNA expression observed in T2D islets may contribute to impaired insulin secretion and disrupted glucose homeostasis.

### 3.3. Suppressing AASS-dependent Lysine Catabolism Reduces Insulin Secretion in Pancreatic Islets and INS1 832/13 ***β*** Cells

To investigate whether the insulinotropic effect of lysine depends on lysine catabolism, we genetically inhibited AASS-dependent lysine catabolism in INS1 832/13 cells and assessed the effects of lysine on insulin secretion. The expression of *Aass* was silenced in INS1 832/13 cells, and reduced expression was confirmed by qPCR analysis (Fig. S2A). We then evaluated the effects of lysine on GSIS in control (NC) versus *Aass*-knockdown (*Aass*-KD) cells. While lysine increased insulin secretion in the presence of high glucose (HG) in both NC and *Aass*-KD cells (Fig. 3A), the stimulatory index (fold change of insulin secretion at HG vs. LG) in the presence of lysine increased only in NC cells and not in *Aass*-KD cells (SI in NC cells: 3.5±0.8 (wo lysine) vs. 4.4±0.8 (with lysine), p-value: 0.0017) (Fig. 3B).

**Figure 3.**
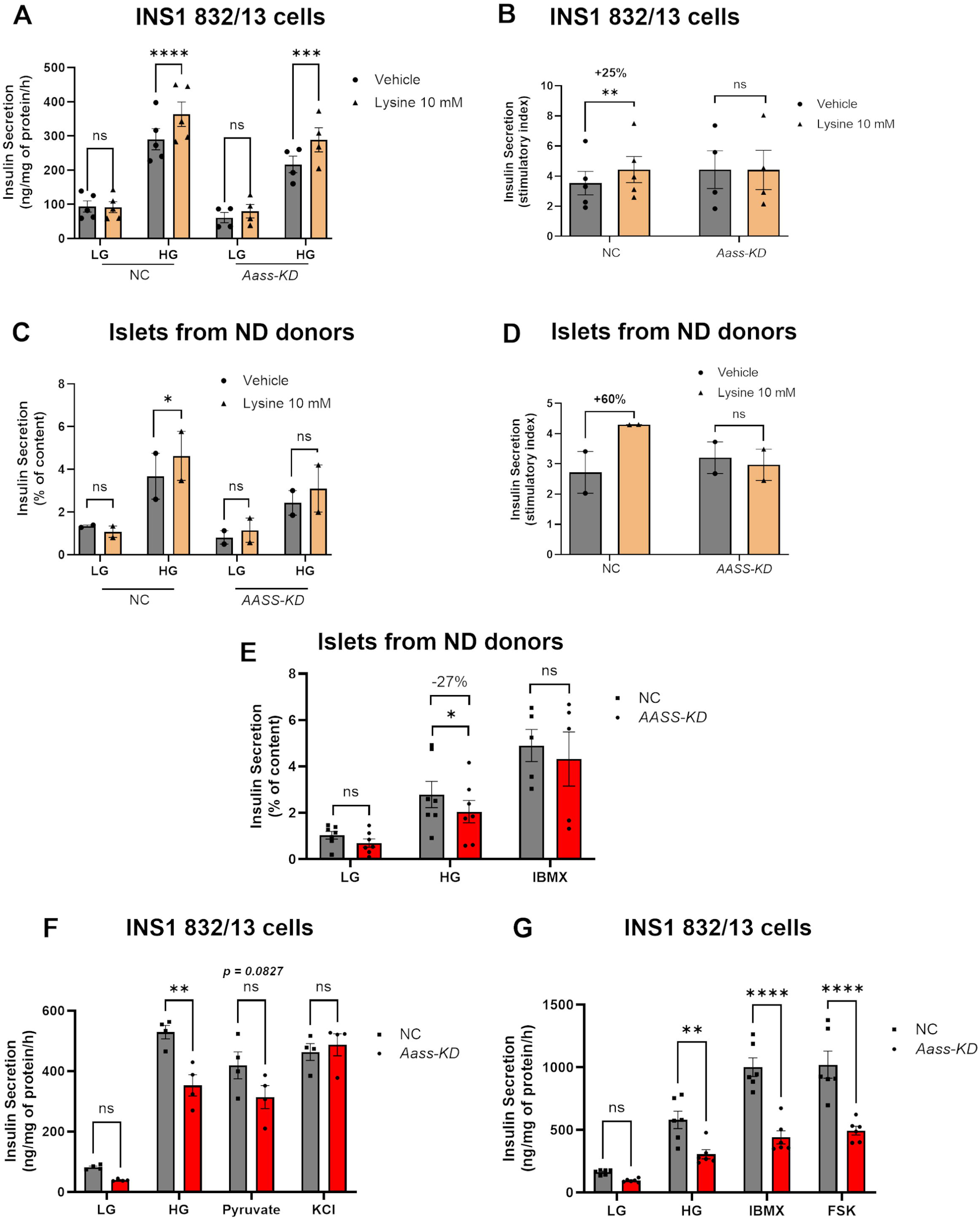
Suppressing AASS-dependent Lysine Catabolism Reduces Insulin Secretion in Pancreatic Islets and INS1 832/13 β Cells. **A, B.** Insulin secretion at LG and HG in the absence and presence of lysine (10 mM) in normal control (NC) and *Aass-KD* INS1 832/13 β cells (n=5). **D.** Calculated stimulatory index (insulin secretion at HG/LG) in presence vs absence of lysine in NC and *Aass-KD* INS1 832/13 β cells (n=5). **C, D.** Insulin secretion in human islets from non-diabetic donors (ND) at low glucose (LG, 2.8 mM) and high glucose (HG, 16.7 mM) in the absence and presence of lysine (10 mM) in ND and *AASS-KD* ND islets (n=2). **D.** Calculated stimulatory index (insulin secretion at HG/LG) in presence vs absence of lysine in NC control and *AASS-KD* ND islets (n=2). **E.** Insulin secretion at LG, HG and HG+IBMX (100 µM) in NC control and *AASS-KD* ND islets (n=7). **F.** Insulin secretion at LG, HG, pyruvate (10 mM) and high K^+^ in NC control and *Aass-KD* INS1 832/13 β cells (n=4). **G.** Insulin secretion at LG, HG, HG+IBMX (100 µM) and HG+FSK (10 µM) in NC control and *Aass-KD* INS1 832/13 β cells (n=6). All panels show individual data points and mean ± s.e.m. Student t-test or Two-way ANOVA (depending on number of compared groups); *p<0.05, **p < 0.01, ***p < 0.001, ****p < 0.0001. AASS, Aminoadipate-Semialdehyde Synthase; KD, knockdown; NC, normal control; ND, normal donor; FSK, forskolin; IBMX, isobutyl-1-methylxanthine.

Next, we tested whether suppressing AASS-dependent lysine catabolism eliminated the insulinotropic effect of lysine in human islets from ND donors. Reduced AASS expression in human islets transfected with AASS vs scramble (control) siRNA was confirmed by qPCR (Fig. S2B). We measured insulin secretion at LG and HG in the presence or absence of lysine in NC versus *AASS*-KD islets (Fig. 3C). Lysine potentiated insulin secretion in control NC islets but not in *AASS*-KD islets (Fig. 3C). As a result, the stimulatory index increases in NC islets (∼60%) but not in *AASS*-KD islets (Fig. 3D).

In a set of independent experiments, we examined whether suppressing AASS-dependent lysine catabolism affected GSIS in islets from ND donors. We observed no differences at LG but a significant reduction of insulin secretion in presence of HG following AASS silencing in human islets (Fig. 3E).

The effect of AASS silencing on GSIS was also evaluated in human EndoC-βH1 β cells, where no significant changes were observed (Fig. S2C). This may be attributed to the fetal phenotype of these cells^32^.

To further confirm whether suppressing AASS-dependent lysine catabolism affects GSIS, we assessed the effects of silenced *Aass* on insulin secretion in response to various metabolic and pharmacologic stimuli in INS1 832/13 β cells (Figs. 3F, G). Silencing *Aass* expression did not changed insulin secretion at LG whereas resulted in significantly reduced insulin secretion at HG (529.1±22.1 vs. 353.0±35.2 ng/mg of protein/h, p-value: 0.0016) and borderline decreases in response to pyruvate stimulation (419.2±44.7 vs. 313.9±38.1 ng/mg of protein/h, p-value: 0.082), but not in response to plasma membrane depolarization with high K^+^ (Fig. 3F). These data suggest that AASS-dependent lysine catabolism contributes to insulin secretion via a mechanism upstream of plasma membrane depolarization likely involving mitochondrial energy metabolism.

The incretin hormone GLP1 amplifies GSIS by promoting cAMP accumulation upon activation of GLP1 receptor^33^. INS1 832/13 cells respond poorly to GLP1 or GLP1R agonists such as Exendin-4^19,34^. Thus, to mimic GLP1R activation, IBMX and forskolin (FSK) were used to trigger cAMP accumulation. IBMX and FSK amplified GSIS in NC cells, but this effect was significantly reduced in *Aass*-KD cells (for IBMX: 1000.9±74.6 vs. 440.5±52.3 ng/mg of protein/h, p-value<0.0001, for FSK: 1021.0±108.1 vs. 494.2±35.1 ng/mg of protein/h, p-value<0.0001) (Fig. 3G).

Overall, these data indicate that an active AASS-dependent catabolism of the endogenous lysine contributes to potentiates GSIS, as well as to the amplification of insulin secretion in response to incretin receptor signalling that culminates in accumulation of cAMP in β cells.

### 3.4. Transcriptomics and Metabolomics Analysis Suggest a Link Between AASS-Dependent Lysine Metabolism and GABA Production/Signaling in ***β*** Cells

To explore the molecular mechanisms by which AASS-dependent lysine catabolism potentiates GSIS, we performed bulk RNA sequencing (RNAseq) and mass-spectrometry-based metabolomics profiling of NC and *Aass*-KD INS1 832/13 cells (Fig. 4).

**Figure 4.**
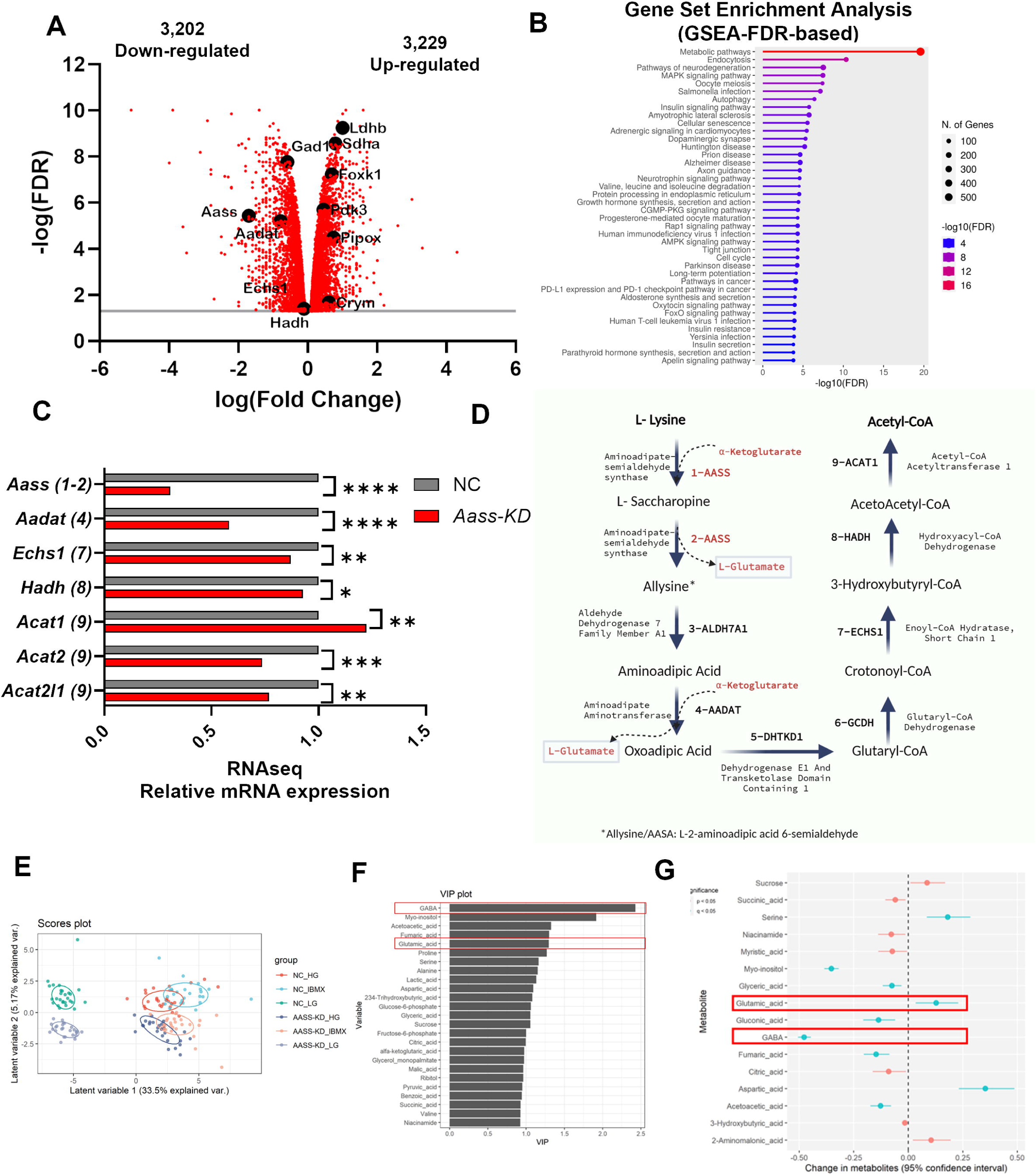
Transcriptomics and Metabolomics Analysis Suggest a Link Between AASS-Dependent Lysine Metabolism and GABA Production/Signaling in β Cells. **A.** Volcano plot and **B.** GSEA based on data from RNAseq analysis of NC and *Aass*-KD INS1 832/13 β cells (n=4). **C.** Expression levels of lysine catabolism pathway-related genes in NC control and *Aass*-KD INS1 832/13 β cells (n=4). **D.** Chart of the lysine catabolism pathway. **E.** PCA score plot showing first two components. **F.** VIP from the predictive component of the OPLS-DA model. **G.** Multivariate ANOVA plot based on data from metabolomics analysis of NC and *Aass*-KD INS1 832/13 β cells (n=7). *p<0.05, **p < 0.01, ***p < 0.001, ****p < 0.0001. GSEA, Gene set enrichment analysis; KD, knockdown; NC, normal control; PCA, principal component analysis; OPLS-DA, orthogonal partial least squares discriminant analysis.

The RNAseq analysis revealed that inhibiting AASS-dependent lysine catabolism significantly impacted global gene expression, with 6,431 differentially expressed genes (DEGs) in *Aass*-KD vs. NC cells. Of these, 3,202 DEGs were downregulated, while 3,229 were upregulated (Fig. 4A). Gene set enrichment analysis (GSEA) highlighted the profound effect of blocking AASS-dependent degradation of the endogenous lysine on overall β cell metabolism (the pathway with the highest false discovery rate (FDR)). Other pathways significantly affected in *Aass*-KD vs. NC cells included neurodegeneration, endocytosis, Alzheimer’s disease, branched-chain amino acid metabolism, autophagy, insulin signaling, long-term potentiation, and insulin secretion, all of which are relevant to β cell physiology (Fig. 4B).

As expected, the RNAseq analysis confirmed a reduction (∼70%) in *Aass* expression in *Aass*-KD cells (Fig. 4C). Supporting the idea that silencing *Aass* resulted in inhibition of the lysine catabolism pathway^17^, the silencing of *Aass*, led to reduced expression of downstream genes in the lysine degradation pathway in *Aass*-KD cells, including *Aadat* (step 4), *Echs1* (step 7), *Hadh* (step 8), and *Acat2* (step 9) (Fig. 4C, D).

To evaluate whether inhibiting lysine degradation at downstream steps also affects insulin secretion, we focused on DHTKD1, the enzyme carrying over the step 5 of lysine degradation pathway (Fig. 4D), and whose expression is also downregulated in T2D islets (Fig. 2D). Thus, we silenced the expression of *Dhtkd1* in INS1 832/13 cells and tested its effect on GSIS. We observed a nominal (p-value: 0.07) decrease in GSIS (Fig. S2D, E).

To assess the relevance of the genetic drift observed upon silencing *Aass* to β cell dysfunction in T2D, we examined how many DEGs in *Aass*-KD cells overlapped with the 395 DEGs we recently reported in pancreatic islets from T2D donors^31^. Remarkably, 33% (129 genes) of the DEGs in T2D islets were also DEGs in *Aass*-KD vs. control cells (Fig. S2F). Several of these overlapping DEGs are known regulators of β cell physiology and T2D pathogenesis (Fig. S2G, highlighted in bold)^34,35^, underscoring the importance of AASS-dependent catabolism of endogenous lysine for β cell function and dysfunction in T2D.

We also conducted mass spectrometry-based metabolomics analysis upon silencing of *Aass* in INS1 832/13 cells (*Aass*-KD vs. NC cells) in response to glucose and glucose+IBMX stimulation. As expected, principal component analysis (PCA) showed clear separation of metabolic clusters based on glucose stimulation (HG vs. LG) and glucose+IBMX, with a less evident separation based on genotype (*Aass*-KD vs. NC cells; Fig. 4E). OPLS-DA was then used to identify the metabolites driving variance between *Aass*-KD and NC cells (Fig. 4F). Strikingly, GABA and glutamate emerged as top metabolites explaining the variation between genotypes. This suggests that suppression of AASS-dependent lysine catabolism may impact a metabolic pathway involving glutamate and GABA, which could account for the loss of GSIS potentiation. Other significant metabolites related to glycolysis (lactate, pyruvate) and the TCA cycle (fumarate, malate, α-ketoglutarate, aspartate, citrate, and succinate) also drove variation between *Aass*-KD and NC cells (Fig. 4F). Multivariate ANOVA confirmed elevated glutamate content and reduced GABA content in *Aass*-KD vs. NC cells (Fig. 4G).

Again, supporting the inhibition of catabolism of endogenous lysine when *Aass* was silenced in *Aass*-KD cells, lysine accumulation was evidenced in *Aass*-KD cells (Fig. S2H). However, the saccharopine pathway (AASS-dependent) in the mitochondrial matrix is not the only route for lysine degradation. The alternative pipecolic pathway, which occurs in the cytosol and peroxisome^17^, may buffer the excess lysine accumulation when AASS-dependent catabolism is suppressed (see pathway chart in Fig. S2I). We therefore examined the expression of genes in the alternative pipecolic pathway and observed an upregulation of *Crym/KR* and *Pipox*, which encode enzymes for steps 2 and 3 of the pipecolic pathway (step 1 gene is unknown), in *Aass*-KD vs. NC cells (Fig. S2J).

To further investigate the relevance of lysine catabolism for GSIS and whether the pipecolic pathway also contributes to GSIS, we silenced *Aass* alone, *Crym/KR* alone, or both *Aass and Crym/KR* in INS1 832/13 cells. Indeed, silencing each pathway individually reduced GSIS, but simultaneous suppression of both pathways led to an additional reduction of GSIS (Fig. S2K).

### 3.5. Silencing of AASS Results in Reduced GABA content, altered GABA Shunt and GABA Signaling

To further explore how AASS-dependent lysine catabolism potentiates insulin secretion in β cells, we investigated the metabolomics findings pointing to GABA and glutamate as key metabolites driving the differences between *Aass*-KD and NC cells. We compared glutamate and GABA cell content in *Aass*-KD and NC cells in response to glucose stimulation (Fig. 5A, B). We observed a trend towards increased glutamate levels (Fig. 5A), while GABA levels significantly decreased in both basal and glucose-stimulated conditions (Fig. 5B).

**Figure 5.**
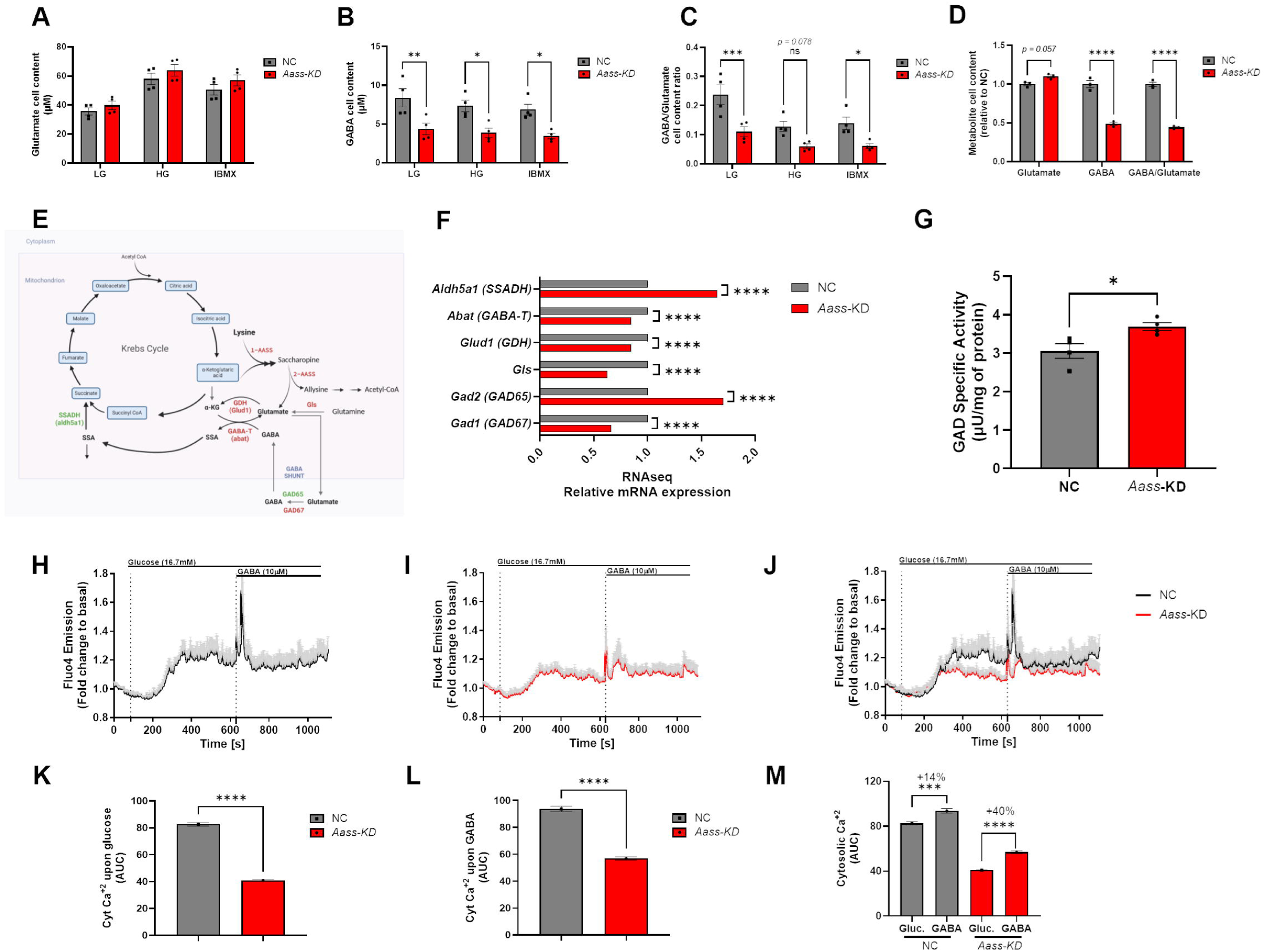
Silencing of AASS Results in Reduced GABA content, altered GABA Shunt and GABA Signaling. **A**. Glutamate cell content, **B**. GABA cell content and **C**. Calculated GABA to Glutamate Cell content ratio from mass spectrometry-based measurement of metabolites in NC control and *Aass*-KD INS1 832/13 β cells (n=4). **D.** Glutamate, GABA and GABA/Glutamate cell content at HG based on targeted metabolomics (n=3). **E.** Chart of GABA shunt pathway, highlighting DEGs in *Aass*-KD vs. NC INS1 832/13 β cells, in green upregulated genes, in red, downregulated genes. **F.** Expression levels of GABA shunt-related genes in NC control and *Aass*-KD INS1 832/13 β cells (n=4). **G.** GAD enzyme activity measured in cell extracts from NC and *Aass*-KD INS1 832/13 β cells (n=4). **H-M.** Average traces of live-cell calcium imaging in response to glucose (16.7 mM) and GABA (10 µM) stimulation in **H.** NC control and **I.** *Aass*-KD INS1 832/13 β cells. **J.** Comparison of the calcium traces in NC vs *Aass*-KD cells (n=6). Average AUC of **K.** glucose-stimulated and **L.** GABA calcium responses and **M.** their comparison in NC vs. *Aass*-KD INS1 832/13 β cells. All panels show individual data points and mean ± s.e.m. Student t-test or Two-way ANOVA (depending on number of compared groups); *p<0.05, **p < 0.01, ***p < 0.001, ****p < 0.0001. GABA, γ-aminobutyric acid; GAD, glutamic acid decarboxylase; KD, knockdown; NC, normal control.

Since glutamate can be converted into GABA in a single step by the enzyme glutamate decarboxylase (GAD), we calculated the GABA-to-glutamate ratio. Interestingly, in control β cells, the GABA-to-glutamate ratio decreased in response to glucose stimulation (Fig. 5C), which aligns with previous studies showing that glucose and others secretagogues stimulates glutamate production^36^ and GABA catabolism to enhance insulin secretion^37^. In *Aass*-KD cells, however, we observed a pronounced reduction in the GABA-to-glutamate ratio at basal and upon glucose stimulation (Fig. 5C). In a set of independent targeted metabolomics experiments, we further confirmed reduced GABA content and GABA-to-glutamate ratio at HG in *Aass*-KD cells (Fig. 5D). These data suggest that the catabolism of endogenous lysine supplies glutamate for its conversion to GABA, regulates the activity of enzymes such as GAD to promote glutamate-to-GABA conversion, or both mechanisms act synergistically. Consequently, suppression of lysine degradation in *Aass*-KD cells leads to reduced GABA production due to decreased glutamate availability or diminished GAD activity. Alternatively, the suppression of lysine degradation in *Aass*-KD cells leads to an accelerated GABA catabolism via GABA shunt, reducing the GABA cell content.

One important metabolic pathway involving both mitochondrial glutamate and GABA is the GABA shunt. In this pathway, α-ketoglutarate (α-KG) from the TCA cycle is transaminated with GABA via GABA transaminase (GABA-T) to produce glutamate and succinic semialdehyde (SSA) in the mitochondrial matrix. Glutamate can then exit the matrix to enter the cytosol, where it can be converted back into GABA via GAD^37,38^ (see scheme in Fig. 5E).

Alongside the marked reduction in the GABA-to-glutamate ratio in *Aass*-KD cells, we observed altered expression of several genes encoding components of the GABA shunt pathway, including *gls* (Glutaminase), *abat* (GABA-T), *glud1* (Glutamate Dehydrogenase 1, GDH), and *aldh5a1* (SSADH), as well as *gad1* (GAD67) and *gad2* (GAD65) (Fig. 5F). Interestingly, in line with this reduced expression of *gls* and *abat* in *Aass*-KD cells, we observed a positive correlation between the expression of *AASS* and *ABAT* (Fig. S3A) and *AASS* and *GLS* (Fig. S3B) in human islets, indicating that the expression of GABA shunt genes may also be regulated by the activity of the AASS-dependent lysine catabolism in human β cells.

To explore whether the GABA shunt pathway is relevant to T2D islet dysfunction, we also examined the expression of GABA shunt genes in pancreatic islets from T2D vs. ND donors. We found downregulation of *ABAT* (Fig. S3C) and upregulation of *GAD1* (*GAD67*) (Fig. S3D) in islets from T2D donors compared to ND donors, supporting the notion that altered GABA shunt metabolism contributes to β cell dysfunction in T2D^38^.

Given the changes in *gad1/gad2* expression and the crucial role of cytosolic GAD in converting glutamate to GABA, we also measured GAD enzyme activity. Interestingly, we found increased GAD activity in *Aass*-KD cells compared to NC cells (Fig. 5G). These findings suggests that the reduced glutamate to GABA conversion when AASS-dependent lysine degradation is suppressed may not result from reduced GAD activity but instead from reduced glutamate production in the mitochondrial matrix and, consequently, from reduced glutamate availability in the cytosol (for GAD). Again, an alternative hypothesis would be an accelerated GABA catabolism via GABA shunt, resulting in reducing GABA cell content.

Another possibility, the observed reduction in the GABA-to-glutamate cell content ratio in *Aass*-KD cells could result from altered rate of secretion of these metabolites from cells, rather than from changes in their metabolism rate. To address this, we examined the expression of subunits of the volume-regulated anion channel (VRAC), recently identified as the GABA efflux channel responsible for non-vesicular GABA release from β cells^39^. We observed altered expression of VRAC subunits, including Lrrc8d (the GABA-specific subunit) (Fig. S3E). However, the extracellular levels of glutamate and GABA mirrored the intracellular levels, with increased glutamate release (Fig. S3F) and decreased GABA release (Fig. S3G), leading to a significantly reduced GABA-to-glutamate ratio in the extracellular medium of *Aass*-KD cells as well (Fig. S3H). Therefore, the differences in extracellular levels are driven by intracellular metabolic changes, not altered rates of release, supporting the conclusion that AASS-dependent lysine degradation modulates glutamate/GABA metabolism.

Supporting a role of AASS-mediated lysine metabolism in GABA signaling, we found altered expression of several genes related to GABA signaling pathways, including genes encoding GABA receptor subunits (Fig. S3I), GABAergic transmission-related proteins (Fig. S3J), and adenylate cyclase subunits (Fig. S3K).

Lastly, we investigated how inhibition of AASS-dependent lysine degradation affects glucose- and GABA-provoked calcium responses. In control cells, glucose induced cytosolic calcium oscillations (Fig. 5I), but, consistent with reduced GSIS, this response was attenuated in *Aass*-KD cells (Fig. 5J-K). When GABA (10 µM) was added to glucose-stimulated NC cells, a ∼14% increase in cytosolic calcium response was observed (Fig. 5L-N). In *Aass*-KD cells, the GABA-induced calcium response was also attenuated (Fig. 5K). However, the relative increases of GABA-induced calcium responses in *Aass*-KD vs. NC cells was elevated with a ∼40% increase in AUC, compared to 14% in NC cells (Fig. 5N). These results suggest that reduced GABA production in *Aass*-KD cells contributes to decreased calcium oscillations and insulin secretion. Nevertheless, in line with the changes in expression of GABA-A receptor subunits in *Aass*-KD cells (Fig. S3I), these cells may adapt to lower extracellular GABA levels becoming more sensitive to GABA signaling.

### 3.6 Exogenous GABA Restores GSIS in Aass-KD Cells

To further investigate the metabolic alterations underlying the reduced GABA/glutamate ratio and impaired GSIS in *Aass*-KD cells, we conducted a systematic functional analysis using pharmacological agents targeting key enzymes in the GABA shunt and GABA signaling pathway (see scheme in Fig. 6A). Specifically, we assessed the effects of allylglycine (GAD inhibitor), vigabatrin (γ-vinyl GABA, GABA-T inhibitor), α-ketoisocaproic acid (α-KIC, GDH activator), leucine + glutamine (GDH activator and glutamate precursor), muscimol (GABA-A receptor agonist), and GABA on insulin secretion.

**Figure 6.**
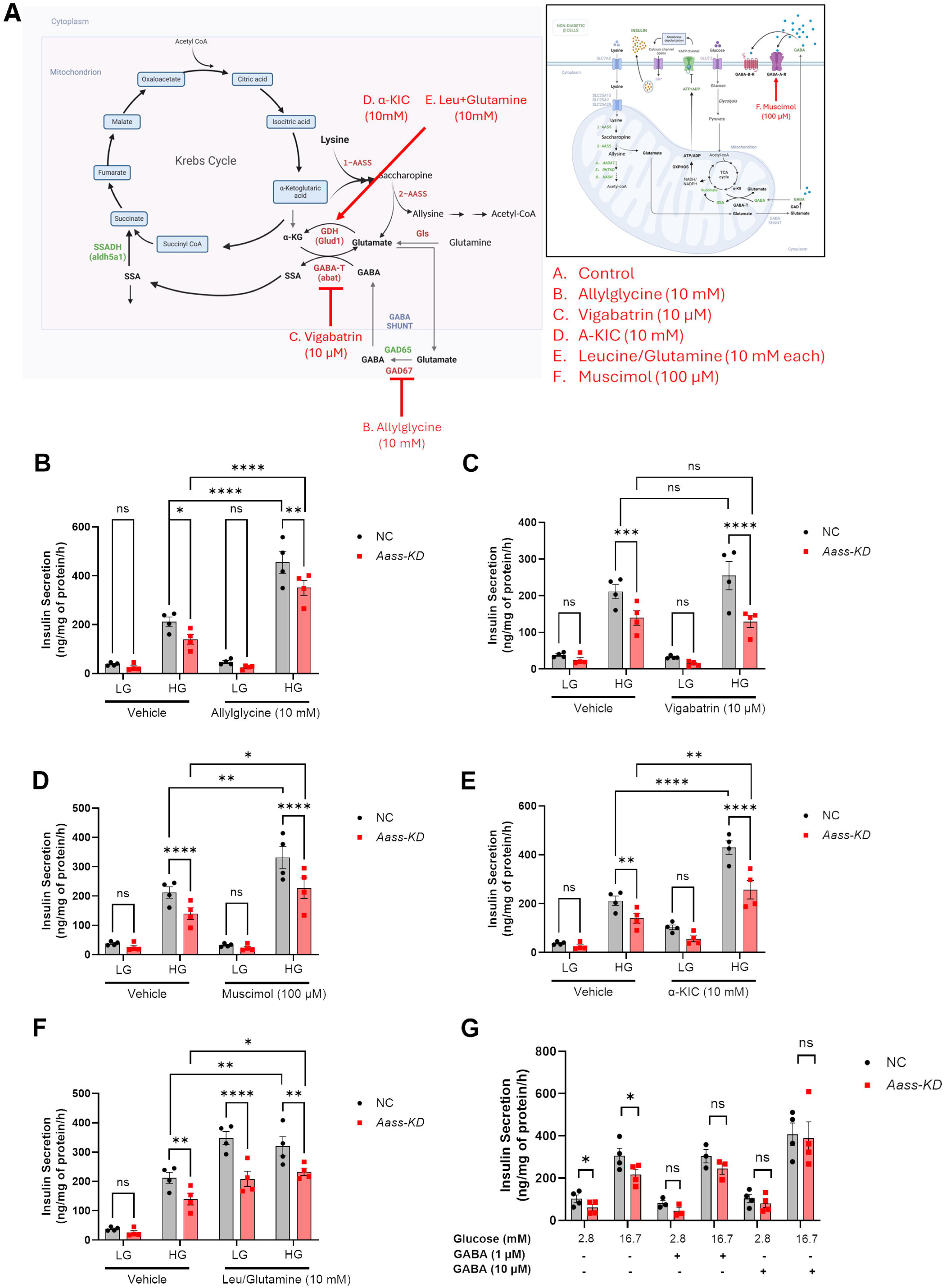
Pharmacological manipulation of GABA shunt and GABA-A receptor signaling and GABA rescue *in Aass-KD Cells*. **A.** Chart of interconnection of AASS-dependent lysine catabolism, TCA cycle, GDH anaplerosis, GABA shunt and GABA-A receptor pathways and specification of pharmacological agents used to manipulates these pathways. **B-G.** Effects on GSIS in NC and *Aass-KD* cells in response to acute manipulation of above mentioned pathways by using the next pharmacological agents: **B.** allylglycine (GAD inhibitor), **C.** vigabatrin (γ-vinyl GABA, GABA-T inhibitor), **D.** α-ketoisocaproic acid (α-KIC, GDH activator), **E.** leucine + glutamine (GDH activator and glutamate precursor), **F.** muscimol (GABA-A receptor agonist), and **G.** GABA (n=4), two way, highlighting DEGs in *Aass*-KD vs. NC INS1 832/13 β cells, in green upregulated genes, in red, downregulated genes.**G.** Insulin secretion at LG and HG in the absence or presence of GABA (1 and 10 µM) in NC control and *Aass*-KD INS1 832/13 β cells (n=4), Two-way ANOVA.

Given the reduced GABA/glutamate ratio in *Aass*-KD cells, we hypothesized that impaired conversion of glutamate to GABA via GAD may be responsible. The previous observations of a change in expression of *gad1/gad2* and increased GAD-specific activity in *Aass*-KD cells also support the participation of this pathway in the *Aass*-KD cells phenotype. To test whether an altered rate of conversion of glutamate to GABA by GAD contributes to the metabolic imbalance and insulin secretory dysfunction in *Aass*-KD cells, we inhibited GAD with allylglycine and measured GSIS (Fig. 6B). Consistent with previous findings in human islets^39^, allylglycine treatment strongly enhanced GSIS in NC cells (+134%). Notably, the increase in GSIS was more pronounced in *Aass*-KD cells (+200%), yet insulin secretion remained lower than in NC cells.

To further investigate the role of GABA catabolism, we focused on GABA transaminase (GABA-T), the enzyme responsible for converting GABA into SSA and α-KG into glutamate. Notably, expression of the gene encoding GABA-T, *Abat*, was downregulated in *Aass*-KD cells. Here, we inhibited GABA-T with vigabatrin and measured GSIS. Vigabatrin had no significant effect on GSIS in either NC or *Aass*-KD cells, suggesting that altered GABA-T activity does not fully explain the reduced GSIS in *Aass*-KD cells (Fig. 6C).

Given the interplay between AASS-dependent lysine catabolism, the GABA shunt, and GDH activity (see scheme in Fig. 6A), we examined whether modulating GDH could provide insights into the metabolic alterations in *Aass*-KD cells. Activation of GDH with α-KIC significantly increased GSIS in NC cells (+90%), but this effect was attenuated in *Aass*-KD cells (+55%) (Fig. 6D). Interestingly, co-stimulation with leucine and glutamine (a glutamate precursor) resulted in a more pronounced increase in GSIS in *Aass*-KD cells compared to NC cells (Fig. 6E). This finding supports the notion that altered GDH activity may be associated with a limited supply of glutamate derived from AASS-dependent lysine catabolism, contributing to the reduced GSIS observed in *Aass*-KD cells.

Since GABA can act as an autocrine excitatory signal through GABA-A receptors to enhance insulin secretion in human β-cells^40^, we next investigated whether GABA-A receptor activation could rescue GSIS in *Aass*-KD cells. Indeed, muscimol treatment increased GSIS in NC cells (+58%), with a more pronounced effect in *Aass*-KD cells (+100%) (Fig. 6F), consistent with the heightened calcium response sensitivity observed in *Aass*-KD cells (Fig. 5M). However, GSIS remained lower in *Aass*-KD compared to NC cells, indicating that GABA-A receptor activation only partially restored insulin secretion.

Finally, we tested whether direct GABA replenishment, which could both activate GABA-A receptors and compensate for potential metabolic imbalances, could restore GSIS in *Aass*-KD cells. At 1 µM GABA, GSIS was partially restored, while at 10 µM GABA, GSIS was fully recovered (Fig. 6G). To explore the clinical relevance of these findings, we assessed the effects of acute GABA stimulation on GSIS in human islets from ND and T2D donors. In ND islets, GABA reduced insulin secretion at low glucose levels (Fig. S3L) and tended to increase the GSIS stimulatory index (Fig. S3M, *p = 0.09*). However, this effect was absent in T2D islets, leading to a reduced stimulatory index (Fig. S3N–O, *p = 0.08*).

### 3.7. Silencing of Aass Results in Altered Glycolysis, Mitochondrial TCA Cycle and ATP production

Mitochondrial GABA metabolism via the GABA shunt pathway plays a crucial role in GSIS by providing anaplerotic substrates to the TCA cycle in β cells^37,38^. In line with this, transcriptomic and metabolomic analyses of *Aass*-KD and NC INS1 832/13 cells also revealed alterations in the expression of genes and the abundance of metabolites related to glycolysis, TCA cycle and oxidative phosphorylation (OXPHOS) (Fig. 7).

**Figure 7.**
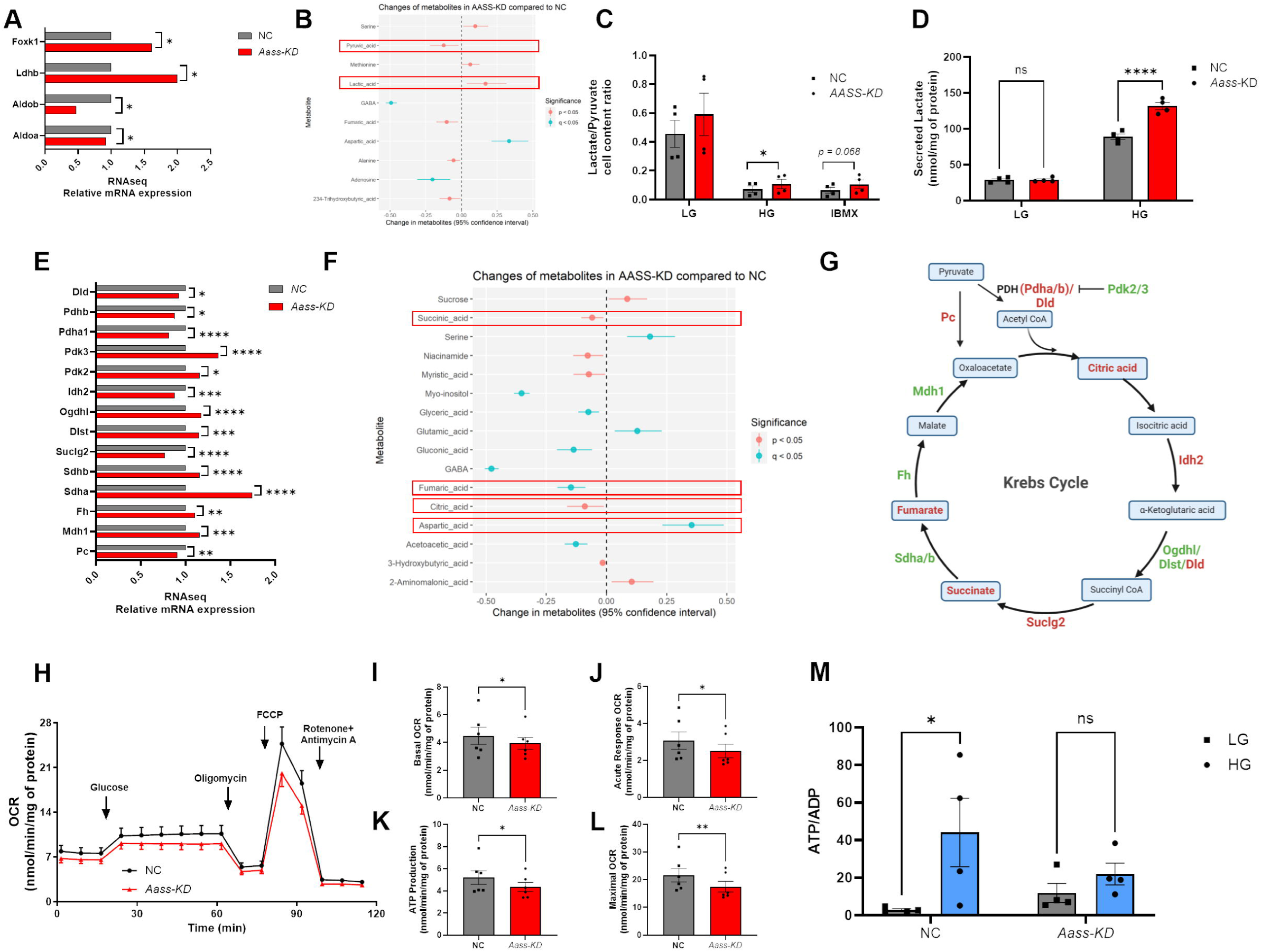
Silencing of *Aass* Results in Altered Glycolysis, Mitochondrial TCA Cycle and ATP production. **A**. Average expression levels of glycolysis-related genes in NC and *Aass*-KD INS1 832/13 β cells (n=4). **B.** Plot showing differences in glycolysis-related metabolites cell contents in NC vs. *Aass*-KD INS1 832/13 β cells (n=7). **C.** Lactate to pyruvate cell content ratio at LG, HG and HG+IBMX in NC control and *Aass*-KD INS1 832/13 β cells (n=4). **D.** Lactate concentration in extracellular medium at LG and HG in NC control vs. *Aass*-KD INS1 832/13 β cells (n=4). **E**. Average expression levels of TCA cycle-related genes in NC control and *Aass*-KD INS1 832/13 β cells (n=4). **F.** Plot showing differences in TCA cycle-related metabolites cell contents in NC control vs. *Aass*-KD INS1 832/13 β cells (n=4). **G.** Chart of TCA cycle pathway, highlighting DEGs and changes in metabolites cell contents in *Aass*-KD vs. NC control INS1 832/13 β cells, in green upregulated genes, in red, downregulated genes or less abundant metabolites. **H-L.** Mitochondrial OCR in NC and *Aass*-KD INS1 832/13 β cells (n=6). **H.** Average OCR traces at LG, HG, Oligomycin (4 mM), FCCP (4 mM) and Antimycin A+Rotenone (1 mM) conditions. **I-L.** Comparison of calculated mitochondrial respiratory parameters in NC vs. *Aass*-KD INS1 832/13 β cells (n=6); **I.** Basal OCR, **J.** Acute Response OCR, **K.** ATP-linked OCR, **L.** Maximal OCR. **M.** Whole-cell ATP to ADP content ratio measured by LC-MS after 60 min of stimulation with LG or HG in NC control vs. *Aass*-KD INS1 832/13 β cells (n=4). All panels show individual data points and mean ± s.e.m. Student t-test or Two-way ANOVA (depending on number of compared groups); *p<0.05, **p < 0.01, ***p < 0.001, ****p < 0.0001. KD, knockdown; NC, normal control; DEG, differentially expressed gene; LC-MS, liquid chromatography-mass spectrometry; OCR, oxygen consumption rate.

We observed increased expression of *Foxk1* and *Ldhb* in *Aass*-KD cells (Fig. 7A). FOXK1/2 (forkhead transcription factors 1/2) are activated during fasting and starvation and promote glycolysis while inhibiting mitochondrial pyruvate oxidation^41^. Consistently, *Aass*-KD cells showed decreased intracellular pyruvate levels and increased lactic acid compared to NC cells (Fig. 7B). These changes suggest impaired mitochondrial glucose metabolism in *Aass*-KD cells, with a shift of pyruvate from entering the TCA cycle to being converted into lactate. This idea is further supported by the elevated lactate-to-pyruvate ratio observed in *Aass*-KD cells in response to glucose (Fig. 7C), along with increased lactate secretion (Fig. 7D).

In support of reduced TCA cycle activity in *Aass*-KD cells, we also observed altered expression of genes encoding mitochondrial TCA cycle proteins (Fig. 7E, G) and reduced levels of key TCA cycle intermediates, including citric acid, succinic acid, and fumaric acid, alongside increased aspartic acid levels (Fig. 7F, G).

The ketone bodies; acetoacetate and 3-hydroxybutyrate serve to anaplerotically feed the TCA cycle and in combination with other substrates stimulates insulin secretion^42^. Coenzyme A conjugated forms of these metabolites are produced along the lysine catabolic pathway (Fig. 4D) and they are significantly reduced in *Aass*-KD compared to NC cells (Fig. 7F).

To further assess how these TCA cycle alterations in *Aass*-KD cells might affect mitochondrial OXPHOS, we measured oxygen consumption rate (OCR) in response to glucose and mitochondrial inhibitors, which allowed us to calculate key respiratory parameters (Fig. 7H-L). The analysis revealed significantly reduced mitochondrial respiration in *Aass*-KD cells compared to NC cells. This included reductions in basal OCR (Fig. 7I), acute response OCR (Fig. 7J), ATP-linked OCR (Fig. 7K), and maximal OCR (Fig. 7L). These findings are consistent with a blunted increase in the ATP/ADP cell content ratio in *Aass*-KD cells following glucose stimulation, whereas control NC cells showed an expected rise (Fig. 7M). These overall reduced TCA cycle and OXPHOS activity in response to suppressing lysine catabolism in *Aass*-KD cells was also aligned with altered expression of multiple genes encoding subunits of Complex I (Fig. S4A), II (Fig. S4B), III (Fig. S4C), IV (Fig. S4D) and V (Fig. S4E) of OXPHOS.

Together, these results indicate that supressing AASS-dependent lysine metabolism in β cells may leads to altered GABA-mediated and/or glutamate-GDH-mediated anaplerosis of the TCA cycle. As a consequence, mitochondrial OXPHOS is impaired and the ATP/ADP ratio reduced, which may contribute to defective insulin secretion.

## 4. Discussion

Lysine is an essential amino acid with known insulinotropic effects *in vivo* in humans and *in vitro* in β cell lines and rodent islets^2,4^. This study investigated whether the insulinotropic action of lysine requires its catabolism via the AASS-dependent pathway and the subsequent production of mitochondrial-derived metabolic coupling factors (MCFs). Our findings demonstrate that AASS-dependent lysine catabolism is crucial for maintaining GABA metabolism and signaling, sustaining mitochondrial function, and potentiating insulin secretion. Consequently, the suppression of this pathway in pancreatic islets from individuals with T2D may contribute to the well-documented depletion of GABA content, impaired GABAergic signaling, and β cell dysfunction during disease progression (graphical abstract in Fig. 8).

**Figure 8.**
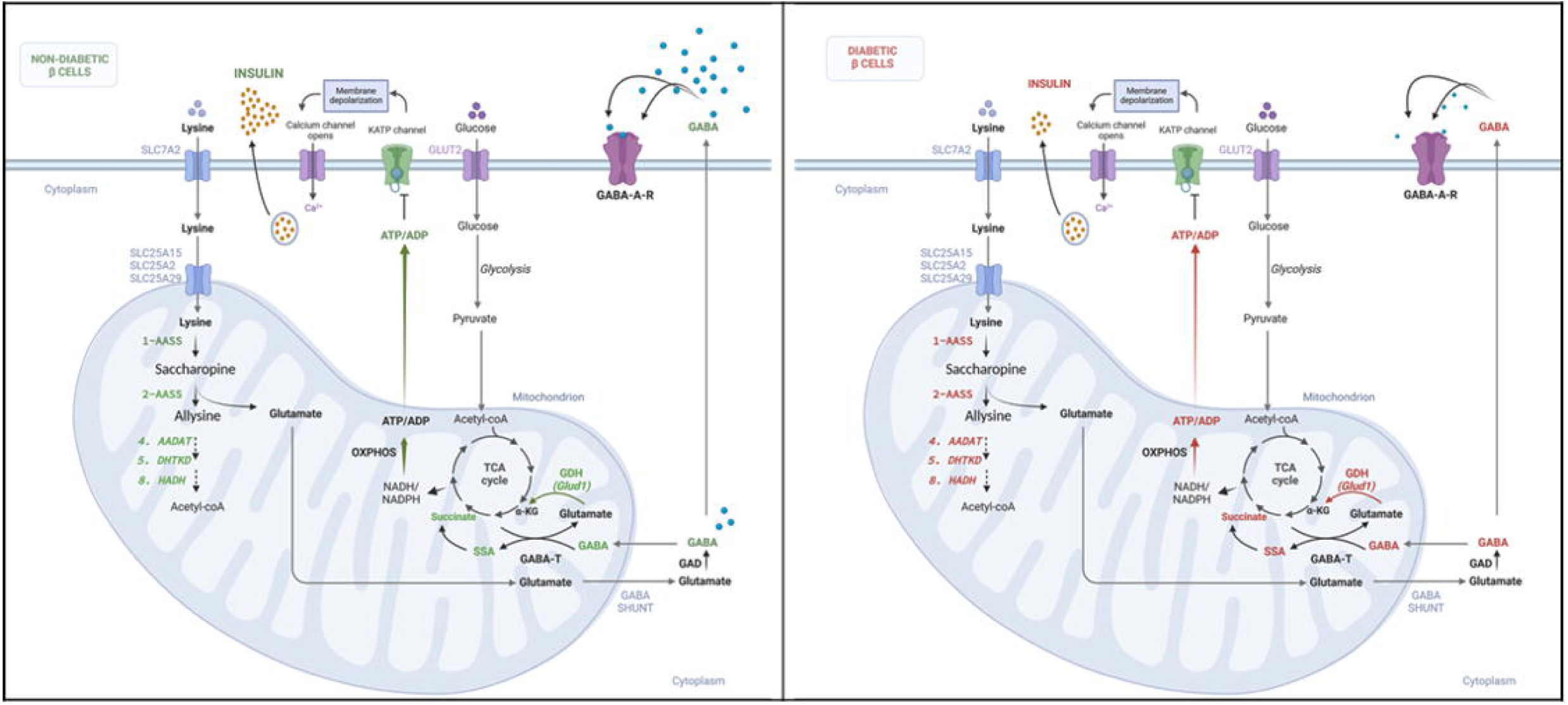
Lysine catabolism potentiates insulin secretion through GABA synthesis in β cells of ND but not of T2D donors. In β cells from non-diabetic (ND) donors, lysine potentiates insulin secretion through its AASS-dependent catabolism. This metabolic pathway provides glutamate, which serves as a substrate for both GABA shunt metabolism and anaplerotic entry into the TCA cycle via GDH, enhancing mitochondrial ATP production and, consequently, insulin secretion. In contrast, in β cells from type 2 diabetes (T2D) donors, the reduced expression of genes involved in lysine catabolism, including AASS, leads to suppressed lysine metabolism. This results in decreased GABA cell content, impaired GABA-A receptor signaling, and diminished anaplerotic fueling of the TCA cycle through the GDH pathway. Consequently, mitochondrial ATP production and insulin secretion are reduced. **Abbreviations:** GABA, γ-aminobutyric acid; GABA-T, GABA transaminase; GAD, glutamic acid decarboxylase; SSA, succinic semialdehyde.

For decades, lysine was thought to exert insulinotropic effects through direct uptake and plasma membrane depolarization, similar to arginine^4^. However, we show that human islets and INS1 832/13 β cells express genes involved in lysine catabolism and that silencing AASS, the rate-limiting enzyme in this pathway, significantly reduces both lysine-stimulated insulin secretion and GSIS. These findings indicate that the insulinotropic effect of lysine is at least partially dependent on its metabolism via AASS, challenging the conventional view of lysine as a mere depolarizing agent.

Our data support the previously underappreciated hypothesis that lysine functions as a metabolic fuel in β cells^43^, consistent with prior metabolomic studies indicating that glucose not only stimulates insulin secretion but also promotes lysine catabolism in β cells^44,45^. Additionally, our finding that lysine potentiates GSIS, at least in part, through its AASS-dependent catabolism aligns with previous reports showing that acute treatment with diazoxide, an ATP-sensitive K□ channel opener that repolarizes the plasma membrane, and verapamil, a blocker of voltage-gated calcium channels (VGCCs), both attenuate lysine’s potentiating effect on GSIS in the rat BRIN-BD11 β-cell line^46^. These results suggest that lysine catabolism contributes to ATP production, facilitating ATP-sensitive K□ channel closure and VGCC opening, ultimately enhancing insulin secretion.

A key finding of this study is the impact of suppressed AASS-dependent lysine catabolism on β cell GABA content. *Aass*-KD cells exhibited a significant reduction in intracellular GABA levels and an altered GABA/glutamate ratio, indicating an imbalance in the metabolic pathways responsible for producing and utilizing these metabolites. Consistent with this, a recent study demonstrated that silencing RHOT1 (MIRO1) in INS1 832/13 cells, mimicking the epigenetically mediated downregulation of its expression in pancreatic islets from T2D donors, resulted in reduced mitochondrial respiration, ATP production, GSIS, and GABA content^47^. Our findings further highlight the functional relevance of reduced GABA levels in *Aass*-KD cells, as exogenous GABA supplementation restored insulin secretion. Given that GABA metabolism via the GABA shunt has been shown to enhance insulin secretion, our results suggest that altered GABA metabolism contributes to the β cell dysfunction observed upon AASS silencing.

Prior studies have yielded conflicting results regarding the effects of GABA transaminase (GABA-T) inhibition on GSIS. While Pizarro-Delgado et al. reported that inhibiting GABA-T with gabaculine reduced GSIS in rat islets^37^, Menegaz et al. found that increasing endogenous GABA by blocking its catabolism (using γ-vinyl-GABA) inhibited pulsatile insulin secretion in human islets^39^. In our β cell model, GABA-T inhibition (with Vigabatrin) did not significantly affect GSIS, suggesting that promotion of the GABA shunt alone is not sufficient to restore insulin secretion in *Aass*-KD cells. Interestingly, we observed that inhibition of GABA biosynthesis using the GAD inhibitor allylglycine enhanced GSIS in control cells, with a stronger effect in *Aass*-KD cells. This suggests that *Aass*-KD cells may have an accelerated GABA biosynthesis rate, potentially as a compensatory response to increased GABA catabolism.

The impaired GSIS observed in *Aass*-KD cells may also stem from disrupted mitochondrial metabolism. Lysine-derived glutamate serves as a substrate for glutamate dehydrogenase (GDH), which fuels the TCA cycle via anaplerosis. Pharmacological activation of GDH with leucine, α-KIC, and 2-aminobicyclo-(2,2,1)-heptane-2-carboxylic acid (BCH) has been shown to enhance GSIS and preserve β cell function, with potential therapeutic applications in T2D^48^. In our study, the reduced GSIS in *Aass*-KD cells correlated with impaired glutamate supply for GDH-mediated anaplerosis, as indicated by the attenuated response to α-KIC and the enhanced GSIS upon co-stimulation with leucine and glutamine.

Beyond GDH-mediated anaplerosis, our transcriptomic and metabolomic analyses revealed widespread metabolic alterations in *Aass*-KD cells, including reduced TCA cycle activity, impaired oxidative phosphorylation (OXPHOS), and a shift from mitochondrial pyruvate oxidation toward lactate production. These disruptions resulted in decreased ATP generation, as reflected by reduced oxygen consumption and a lower ATP/ADP ratio. Since ATP-sensitive K+ channel closure is required for membrane depolarization and calcium influx, both essential for insulin secretion^49^, these findings provide a mechanistic explanation for the insulin secretion defects observed in *Aass*-KD cells.

GABA acts as an autocrine excitatory signal in β cells via GABA-A receptors, modulating calcium oscillations and insulin secretion^40^. Consistently, *Aass*-KD cells exhibited attenuated glucose-stimulated calcium oscillations and reduced GSIS. However, these cells also showed an increased sensitivity to GABA-induced calcium responses, along with upregulation of GABA-A receptor subunits, suggesting a compensatory adaptation to reduced extracellular GABA levels.

In human islets, we observed that acute GABA exposure decreased insulin secretion at low glucose concentrations, while tending to increase GSIS. This effect was absent in T2D islets, potentially due to disrupted GABA signaling. Notably, the reduced expression of lysine catabolic genes in T2D donor islets, together with the marked depletion of GABA in *Aass*-KD cells, suggests that impaired AASS-dependent lysine catabolism may contribute to the GABA deficiency observed in β cells from individuals with T2D^13,39,50,51^. This aligns with prior findings showing that genetic variants in GAD1, a key gene in GABA synthesis, are associated with T2D risk^52^. Additionally, the expression of several genes related to GABA synthesis (i.e.: GAD1) and signalling (i.e.: GABRA1/GABRA2) is altered in pancreatic islets of T2D donors^31^.

Lysine catabolism is known to generate ketone bodies in tissues such as the liver, kidney, and nervous system^53–55^. In β cells, we observed a reduction in ketone bodies (acetoacetate and 3-hydroxybutyrate) upon AASS silencing, suggesting that lysine-derived ketones may contribute to β cell anaplerosis. Given that ketone bodies can serve as alternative metabolic fuels to support TCA cycle activity and ATP production^42^, their depletion may further exacerbate the metabolic dysfunction and impaired insulin secretion observed in *Aass*-KD cells.

## 5. Conclusion

Our findings highlight AASS-dependent lysine catabolism as a crucial regulator of β cell function, supporting both mitochondrial metabolism and GABAergic signaling. The suppression of this pathway leads to reduced GABA cell content, disrupted mitochondrial energy metabolism, impaired calcium dynamics, and defective insulin secretion. The observed reduction in AASS expression and lysine catabolic genes in T2D donor islets suggests that dysfunction in this pathway may contribute to β cell GABA depletion and metabolic dysfunction in diabetes (graphical abstract in Fig. 8).

Therapeutically, strategies that restore lysine catabolism or enhance GABA content and signaling may hold potential for preserving β cell function in T2D. Future studies should explore whether targeting this pathway could improve insulin secretion and β cell resilience in diabetes.

## Authorship contributions

L.R.C. conceptualized and designed the study. F.M, L.R.C., Q.G., M.M., K.T., A.L. performed experiments and analyzed data. Q.G., T.M. performed statistical data analyses and assisted in experimental design. L.R.C., H.M., M.F., T.M. contributed to interpretation of data. L.R.C. wrote the manuscript. All authors revised the manuscript and approved the final version to be published. L.R.C. is the guarantor of this work and, as such, had full access to all the data in the study and takes responsibility for the integrity of the data and the accuracy of the data analysis.

## Acknowledgements

We thank the Human Tissue Laboratory within EXODIAB/Lund University Diabetes Center. This work includes data and/or analyses from HumanIslets.com funded by the Canadian Institutes of Health Research, JDRF Canada, and Diabetes Canada (5-SRA-2021-1149-S-B/TG 179092) with data from islets isolated by the Alberta Diabetes Institute IsletCore with the support of the Human Organ Procurement and Exchange program, Trillium Gift of Life Network, BC Transplant, Quebec Transplant, and other Canadian organ procurement organizations with written informed donor consent as approved by the Human Research Ethics Board at the University of Alberta (Pro00013094).

We would like to express our gratitude to Professor Patrick Macdonald from the University of Alberta and Jelena Kolic from the University of British Columbia, Canada, for their valuable assistance in clarifying aspects related to the data from humanislets.com. We also thank Anna-Maria Veljanovska Ramsay and Laila Jacobsson from Lund University for their technical support.

We acknowledge Christian Grønbæk and The Single-Cell Omics platform at the Novo Nordisk Foundation Center for Basic Metabolic Research (CBMR) for technical and computational expertise.

## Prior Presentation

Preliminary results of this study were presented at the 58^th^ and 60^th^ Annual Meeting of the EASD, 2022 and 2024, respectively.

## Supplementary Figure Legends

**Figure S1. A.** Table detailing characteristics of the ND and T2D donors of pancreatic islets utilized for experiments in Fig. 1 and 3. **B, C.** Spearman rank correlation analysis of expression levels of genes related to the lysine catabolism pathway in human islets. **B.** Correlation between *AASS* and *ALDH7A1* mRNA levels. **C.** Correlation between *AASS* and *DHTKD1* mRNA levels. **D.** Immunohistochemistry staining for AASS (green) and INSULIN (red) proteins and nuclei (DAPI, blue) in human pancreas sections from ND and T2D donors. **E-G.** Regression analysis between AASS protein levels measured by mass-spectrometry in human islets and; **E.** diabetes status; ND (n=118) vs. T2D (n=16), **F.** β cells proportion and **G.** normalized HbA1c (data obtained from www.humanislets.com)^56^.

**Figure S2. A.** *Aass* mRNA expression quantified by qPCR in NC control vs. *Aass*-KD INS1 832/13 β cells**. B**. *AASS* mRNA expression quantified by qPCR in NC control vs. *AASS*-KD human islets from ND donors. **C.** Insulin secretion at LG, HG and HG+IBMX (100 µM) in NC control and *AASS-KD* EndoC-βH1 cells (n=4). **D.** *Dhtkd1* mRNA expression quantified by qPCR in NC control vs. *Dhtkd1*-KD INS1 832/13 β cells (n=4)**. E.** Insulin secretion at LG and HG in NC control vs. *Dhtkd1*-KD INS1 832/13 β cells (n=4). **F.** Venn Diagram showing the intersection between DEGs in T2D vs. ND islets and DEGs in *Aass*-KD INS1 832/13 β cells. **G.** List of gene names in such intersection. **H.** Lysine cell content measured by mass spectrometry in NC control and *Aass*-KD INS1 832/13 β cells (n=4). **I.** Chart illustrating the two lysine catabolic pathways; the Saccharopine and the Pipecolate pathways. **J.** mRNA expression of the Pipecolate lysine catabolism pathway genes in NC control and *Aass*-KD INS1 832/13 β cells (n=4). **K.** Insulin secretion at LG and HG in NC control, *Aass*-KD, *Crym*-KD or *Aass+Crym*-KD INS1 832/13 β cells (n=6). All panels show individual data points and mean ± s.e.m. Student t-test or Two-way ANOVA (depending on number of compared groups); *p<0.05, **p < 0.01, ***p < 0.001, ****p < 0.0001.

**Figure S3. A, B.** Spearman rank correlation analysis of expression levels of genes related to the GABA shunt pathway in human islets; **A.** Correlation between *AASS* and *ABAT* mRNA levels. **B.** Correlation between *AASS* and *GLS* mRNA levels. **C, D.** mRNA levels of genes related to the GABA shunt pathway; **C.** *ABAT* and **D.** *GAD1* in human islets from ND (n=160) vs. T2D (n=35) donors. **E.** mRNA expression of VRAC, GABA transporter, subunits genes in NC control and *Aass*-KD INS1 832/13 β cells (n=4). **F-H.** Extracellular concentration measured by mass-spectrometry after 1h stimulation with LG and HG in NC control and *Aass*-KD INS1 832/13 β cells of next metabolites, **F.** Glutamate, **G.** GABA and **H.** GABA to Glutamate ratio (n=4). **I-K.** mRNA expression of the **I.** GABA receptors subunits genes, J. GABAergic transmission-related genes and K. adenylate cyclase subunits genes in NC control and *Aass*-KD INS1 832/13 β cells (n=4). **L-O.** Insulin secretion in human pancreatic islets from **L, M.** ND and **N, O**. T2D donors at LG and HG in the absence and presence of GABA (10 µM) (n=3). **M, N.** Calculated GABA effects on stimulatory index (insulin secretion at HG/LG) in **M.** ND and **O.** T2D islets (n=3). All panels show individual data points and mean ± s.e.m. Student t-test or Two-way ANOVA (depending on number of compared groups); *p<0.05, **p < 0.01, ***p < 0.001, ****p < 0.0001.

**Figure S4. A-E.** mRNA expression of OXPHOS complexes genes; A. Complex I, B. Complex II, C. Complex III, D. Complex IV and E. Complex V in NC control vs. *Aass*-KD INS1 832/13 β cells (n=4). All panels show individual data points and mean ± s.e.m. Student t-test or Two-way ANOVA (depending on number of compared groups); *p<0.05, **p < 0.01, ***p < 0.001, ****p < 0.0001.

## References

1 Kolic, J. et al. Proteomic predictors of individualized nutrient-specific insulin secretion in health and disease. Cell Metab 36, 1619–1633 e1615 (2024). 10.1016/j.cmet.2024.06.001

2 Floyd, J. C., Jr., Fajans, S. S., Conn, J. W., Knopf, R. F. & Rull, J. Stimulation of insulin secretion by amino acids. J Clin Invest 45, 1487–1502 (1966). 10.1172/JCI105456

3 Nilsson, M., Stenberg, M., Frid, A. H., Holst, J. J. & Bjorck, I. M. Glycemia and insulinemia in healthy subjects after lactose-equivalent meals of milk and other food proteins: the role of plasma amino acids and incretins. Am J Clin Nutr 80, 1246–1253 (2004). 10.1093/ajcn/80.5.1246

4 Liu, Z., Jeppesen, P. B., Gregersen, S., Chen, X. & Hermansen, K. Dose- and Glucose-Dependent Effects of Amino Acids on Insulin Secretion from Isolated Mouse Islets and Clonal INS-1E Beta-Cells. Rev Diabet Stud 5, 232–244 (2008). 10.1900/RDS.2008.5.232

5 Newsholme, P., Bender, K., Kiely, A. & Brennan, L. Amino acid metabolism, insulin secretion and diabetes. Biochem Soc Trans 35, 1180–1186 (2007). 10.1042/BST0351180

6 Kolic, J., Sun, W. G., Johnson, J. D. & Guess, N. Amino acid-stimulated insulin secretion: a path forward in type 2 diabetes. Amino Acids 55, 1857–1866 (2023). 10.1007/s00726-023-03352-8

7 Papes, F., Surpili, M. J., Langone, F., Trigo, J. R. & Arruda, P. The essential amino acid lysine acts as precursor of glutamate in the mammalian central nervous system. FEBS Lett 488, 34–38 (2001). 10.1016/s0014-5793(00)02401-7

8 Markovitz, P. J. & Chuang, D. T. The bifunctional aminoadipic semialdehyde synthase in lysine degradation. Separation of reductase and dehydrogenase domains by limited proteolysis and column chromatography. J Biol Chem 262, 9353–9358 (1987).

9 Maechler, P. & Wollheim, C. B. Mitochondrial glutamate acts as a messenger in glucose-induced insulin exocytosis. Nature 402, 685–689 (1999). 10.1038/45280

10 Prentki, M., Matschinsky, F. M. & Madiraju, S. R. Metabolic signaling in fuel-induced insulin secretion. Cell Metab 18, 162–185 (2013). 10.1016/j.cmet.2013.05.018

11 Gheni, G. et al. Glutamate acts as a key signal linking glucose metabolism to incretin/cAMP action to amplify insulin secretion. Cell Rep 9, 661–673 (2014). 10.1016/j.celrep.2014.09.030

12 Baekkeskov, S. et al. Identification of the 64K autoantigen in insulin-dependent diabetes as the GABA-synthesizing enzyme glutamic acid decarboxylase. Nature 347, 151–156 (1990). 10.1038/347151a0

13 Korol, S. V. et al. Functional Characterization of Native, High-Affinity GABA(A) Receptors in Human Pancreatic beta Cells. EBioMedicine 30, 273–282 (2018). 10.1016/j.ebiom.2018.03.014

14 Hagan, D. W., Ferreira, S. M., Santos, G. J. & Phelps, E. A. The role of GABA in islet function. Front Endocrinol (Lausanne) 13, 972115 (2022). 10.3389/fendo.2022.972115

15 Razquin, C. et al. Lysine pathway metabolites and the risk of type 2 diabetes and cardiovascular disease in the PREDIMED study: results from two case-cohort studies. Cardiovasc Diabetol 18, 151 (2019). 10.1186/s12933-019-0958-2

16 Wang, T. J. et al. 2-Aminoadipic acid is a biomarker for diabetes risk. J Clin Invest 123, 4309–4317 (2013). 10.1172/JCI64801

17 Leandro, J. & Houten, S. M. The lysine degradation pathway: Subcellular compartmentalization and enzyme deficiencies. Mol Genet Metab 131, 14–22 (2020). 10.1016/j.ymgme.2020.07.010

18 Asplund, O. et al. Islet Gene View-a tool to facilitate islet research. Life Sci Alliance 5 (2022). 10.26508/lsa.202201376

19 Hohmeier, H. E. et al. Isolation of INS-1-derived cell lines with robust ATP-sensitive K+ channel-dependent and -independent glucose-stimulated insulin secretion. Diabetes 49, 424–430 (2000).

20 Ravassard, P. et al. A genetically engineered human pancreatic beta cell line exhibiting glucose-inducible insulin secretion. J Clin Invest 121, 3589–3597 (2011). 10.1172/JCI58447

21 Taneera, J. et al. Identification of novel genes for glucose metabolism based upon expression pattern in human islets and effect on insulin secretion and glycemia. Hum Mol Genet 24, 1945–1955 (2015). 10.1093/hmg/ddu610

22 Sato, S. et al. Atlas of exercise metabolism reveals time-dependent signatures of metabolic homeostasis. Cell Metab 34, 329–345 e328 (2022). 10.1016/j.cmet.2021.12.016

23 Cataldo, L. R. et al. The human batokine EPDR1 regulates beta-cell metabolism and function. Mol Metab 66, 101629 (2022). 10.1016/j.molmet.2022.101629

24 Ewels, P. A. et al. The nf-core framework for community-curated bioinformatics pipelines. Nat Biotechnol 38, 276–278 (2020). 10.1038/s41587-020-0439-x

25 Frankish, A. et al. GENCODE reference annotation for the human and mouse genomes. Nucleic Acids Res 47, D766–D773 (2019). 10.1093/nar/gky955

26 Robinson, M. D., McCarthy, D. J. & Smyth, G. K. edgeR: a Bioconductor package for differential expression analysis of digital gene expression data. Bioinformatics 26, 139–140 (2010). 10.1093/bioinformatics/btp616

27 The Gene Ontology, C. The Gene Ontology Resource: 20 years and still GOing strong. Nucleic Acids Res 47, D330–D338 (2019). 10.1093/nar/gky1055

28 Fabregat, A. et al. The Reactome Pathway Knowledgebase. Nucleic Acids Res 46, D649–D655 (2018). 10.1093/nar/gkx1132

29 Wu, D. & Smyth, G. K. Camera: a competitive gene set test accounting for inter-gene correlation. Nucleic Acids Res 40, e133 (2012). 10.1093/nar/gks461

30 Johnson, W. E., Li, C. & Rabinovic, A. Adjusting batch effects in microarray expression data using empirical Bayes methods. Biostatistics 8, 118–127 (2007). 10.1093/biostatistics/kxj037

31 Bacos, K. et al. Type 2 diabetes candidate genes, including PAX5, cause impaired insulin secretion in human pancreatic islets. J Clin Invest 133 (2023). 10.1172/JCI163612

32 Lawlor, N. et al. Multiomic Profiling Identifies cis-Regulatory Networks Underlying Human Pancreatic beta Cell Identity and Function. Cell Rep 26, 788–801 e786 (2019). 10.1016/j.celrep.2018.12.083

33 Drucker, D. J. The biology of incretin hormones. Cell Metab 3, 153–165 (2006). 10.1016/j.cmet.2006.01.004

34 Cataldo, L. R. et al. The MafA-target gene PPP1R1A regulates GLP1R-mediated amplification of glucose-stimulated insulin secretion in beta-cells. Metabolism 118, 154734 (2021). 10.1016/j.metabol.2021.154734

35 Gheibi, S. et al. Reduced Expression Level of Protein Phosphatase PPM1E Serves to Maintain Insulin Secretion in Type 2 Diabetes. Diabetes 72, 455–466 (2023). 10.2337/db22-0472

36 Karaca, M., Frigerio, F. & Maechler, P. From pancreatic islets to central nervous system, the importance of glutamate dehydrogenase for the control of energy homeostasis. Neurochem Int 59, 510–517 (2011). 10.1016/j.neuint.2011.03.024

37 Pizarro-Delgado, J., Braun, M., Hernandez-Fisac, I., Martin-Del-Rio, R. & Tamarit-Rodriguez, J. Glucose promotion of GABA metabolism contributes to the stimulation of insulin secretion in beta-cells. Biochem J 431, 381–389 (2010). 10.1042/BJ20100714

38 Tamarit-Rodriguez, J. Metabolic Role of GABA in the Secretory Function of Pancreatic beta-Cells: Its Hypothetical Implication in beta-Cell Degradation in Type 2 Diabetes. Metabolites 13 (2023). 10.3390/metabo13060697

39 Menegaz, D. et al. Mechanism and effects of pulsatile GABA secretion from cytosolic pools in the human beta cell. Nat Metab 1, 1110–1126 (2019). 10.1038/s42255-019-0135-7

40 Braun, M. et al. Gamma-aminobutyric acid (GABA) is an autocrine excitatory transmitter in human pancreatic beta-cells. Diabetes 59, 1694–1701 (2010). 10.2337/db09-0797

41 Sukonina, V. et al. FOXK1 and FOXK2 regulate aerobic glycolysis. Nature 566, 279–283 (2019). 10.1038/s41586-019-0900-5

42 MacDonald, M. J. et al. Acetoacetate and beta-hydroxybutyrate in combination with other metabolites release insulin from INS-1 cells and provide clues about pathways in insulin secretion. Am J Physiol Cell Physiol 294, C442–450 (2008). 10.1152/ajpcell.00368.2007

43 Sener, A. et al. Stimulus-secretion coupling of arginine-induced insulin release: comparison with lysine-induced insulin secretion. Endocrinology 124, 2558–2567 (1989). 10.1210/endo-124-5-2558

44 Lorenz, M. A., El Azzouny, M. A., Kennedy, R. T. & Burant, C. F. Metabolome response to glucose in the beta-cell line INS-1 832/13. J Biol Chem 288, 10923–10935 (2013). 10.1074/jbc.M112.414961

45 Huang, M. & Joseph, J. W. Assessment of the metabolic pathways associated with glucose-stimulated biphasic insulin secretion. Endocrinology 155, 1653–1666 (2014). 10.1210/en.2013-1805

46 McClenaghan, N. H., Barnett, C. R., O’Harte, F. P. & Flatt, P. R. Mechanisms of amino acid-induced insulin secretion from the glucose-responsive BRIN-BD11 pancreatic B-cell line. J Endocrinol 151, 349–357 (1996). 10.1677/joe.0.1510349

47 Ronn, T. et al. Genes with epigenetic alterations in human pancreatic islets impact mitochondrial function, insulin secretion, and type 2 diabetes. Nat Commun 14, 8040 (2023). 10.1038/s41467-023-43719-9

48 Gohring, I. & Mulder, H. Glutamate dehydrogenase, insulin secretion, and type 2 diabetes: a new means to protect the pancreatic beta-cell? J Endocrinol 212, 239–242 (2012). 10.1530/JOE-11-0481

49 Munoz F, F. M., Mulder H, Moritz T and Cataldo LR. Unique β cells metabolic features lost in type 2 diabetes. Acta Physiologica In Press (2024).

50 Weitz, J., Menegaz, D. & Caicedo, A. Deciphering the Complex Communication Networks That Orchestrate Pancreatic Islet Function. Diabetes 70, 17–26 (2021). 10.2337/dbi19-0033

51 Taneera, J. et al. gamma-Aminobutyric acid (GABA) signalling in human pancreatic islets is altered in type 2 diabetes. Diabetologia 55, 1985–1994 (2012). 10.1007/s00125-012-2548-7

52 Suzuki, K. et al. Genetic drivers of heterogeneity in type 2 diabetes pathophysiology. Nature 627, 347–357 (2024). 10.1038/s41586-024-07019-6

53 Papes, F., Kemper, E. L., Cord-Neto, G., Langone, F. & Arruda, P. Lysine degradation through the saccharopine pathway in mammals: involvement of both bifunctional and monofunctional lysine-degrading enzymes in mouse. Biochem J 344 **Pt 2**, 555–563 (1999).

54 Markovitz, P. J., Chuang, D. T. & Cox, R. P. Familial hyperlysinemias. Purification and characterization of the bifunctional aminoadipic semialdehyde synthase with lysine-ketoglutarate reductase and saccharopine dehydrogenase activities. J Biol Chem 259, 11643–11646 (1984).

55 Rao, V. V., Pan, X. & Chang, Y. F. Developmental changes of L-lysine-ketoglutarate reductase in rat brain and liver. Comp Biochem Physiol B 103, 221–224 (1992). 10.1016/0305-0491(92)90435-t

56 Ewald, J. D. et al. HumanIslets.com: Improving accessibility, integration, and usability of human research islet data. Cell Metab (2024). 10.1016/j.cmet.2024.09.001

