## Supplementary figures and images for "Lysine Potentiates Insulin Secretion via AASS-Dependent Catabolism and Regulation of GABA Content and Signaling"

### Figure S1

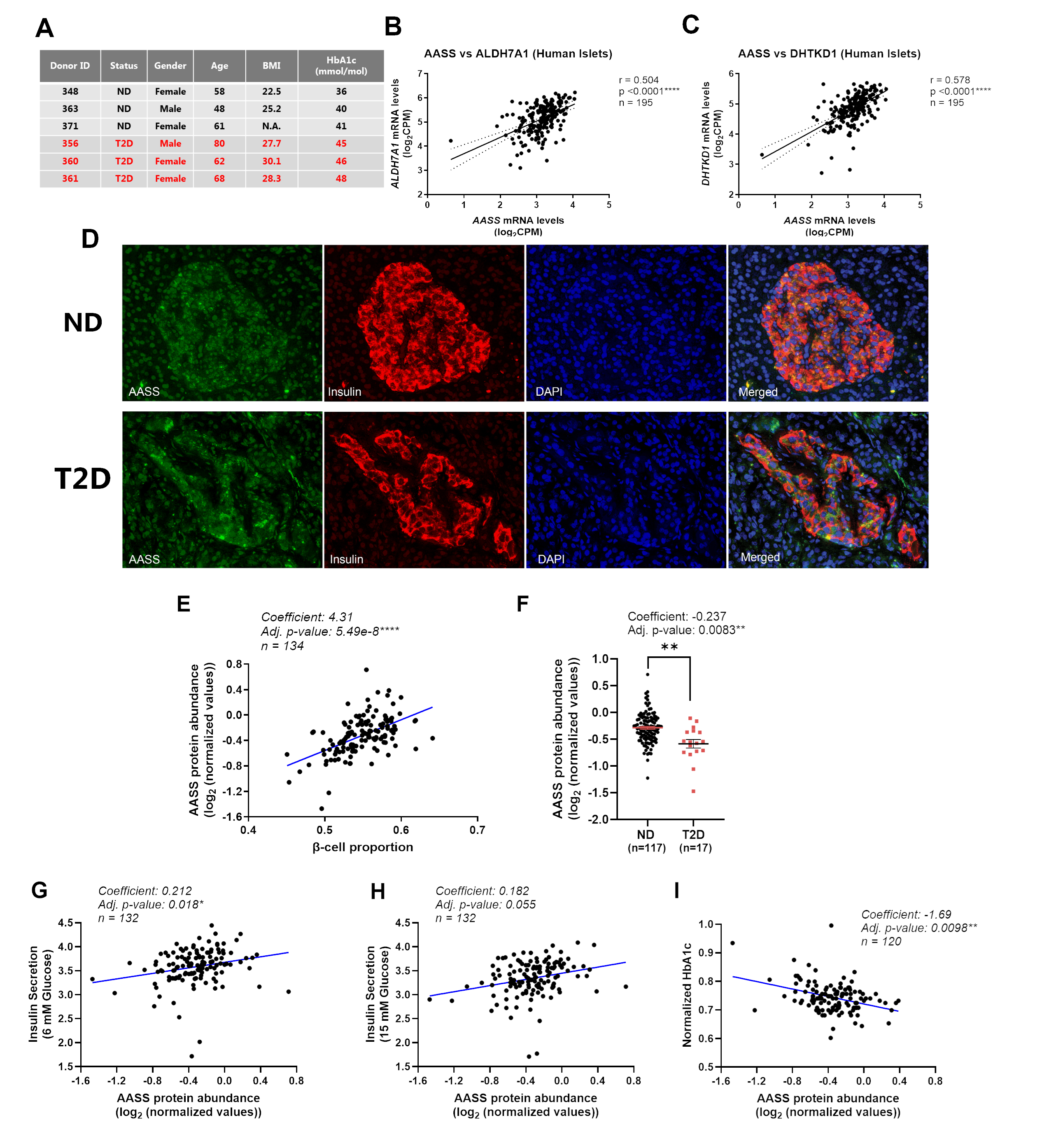

### Figure S2

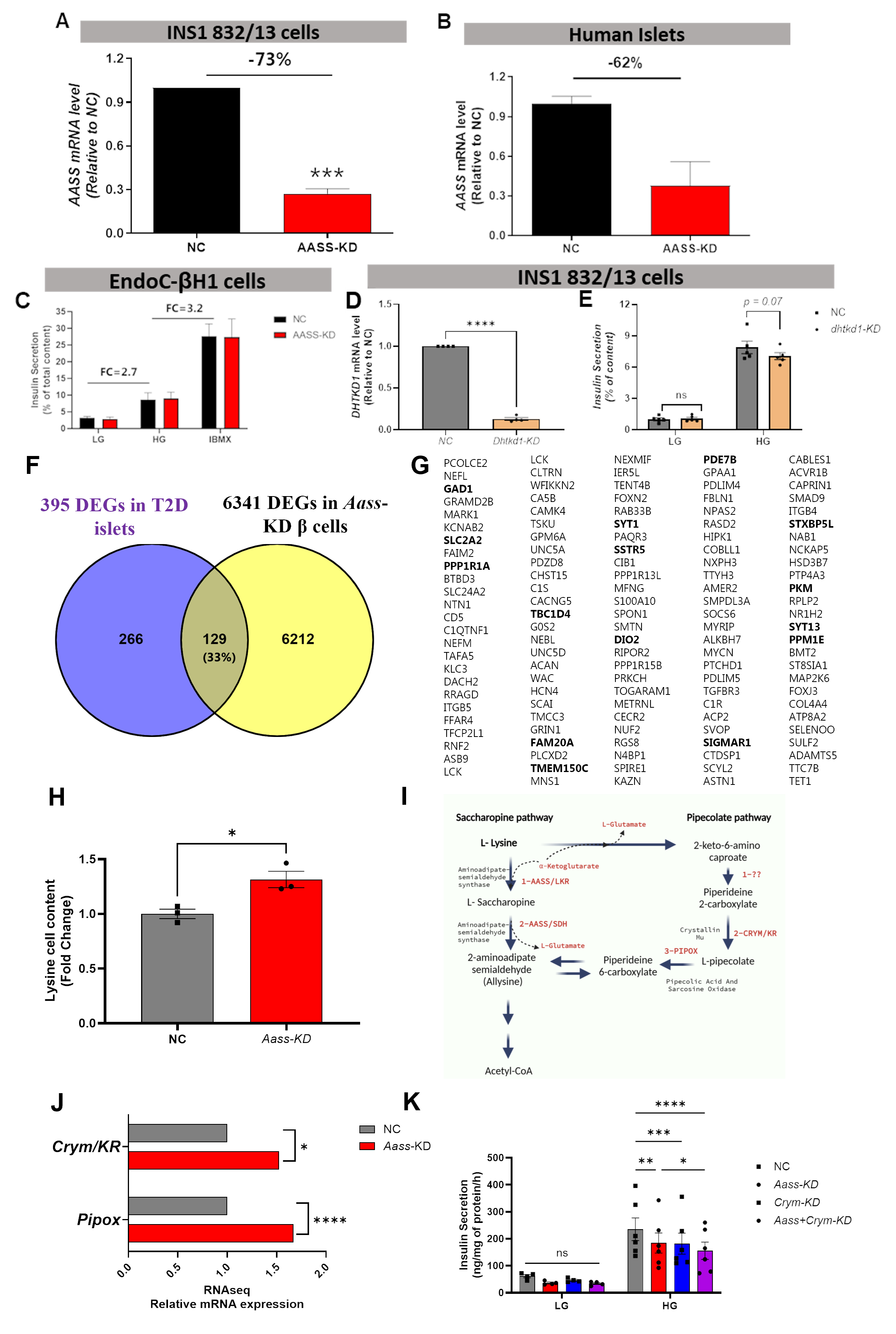

### Figure S3

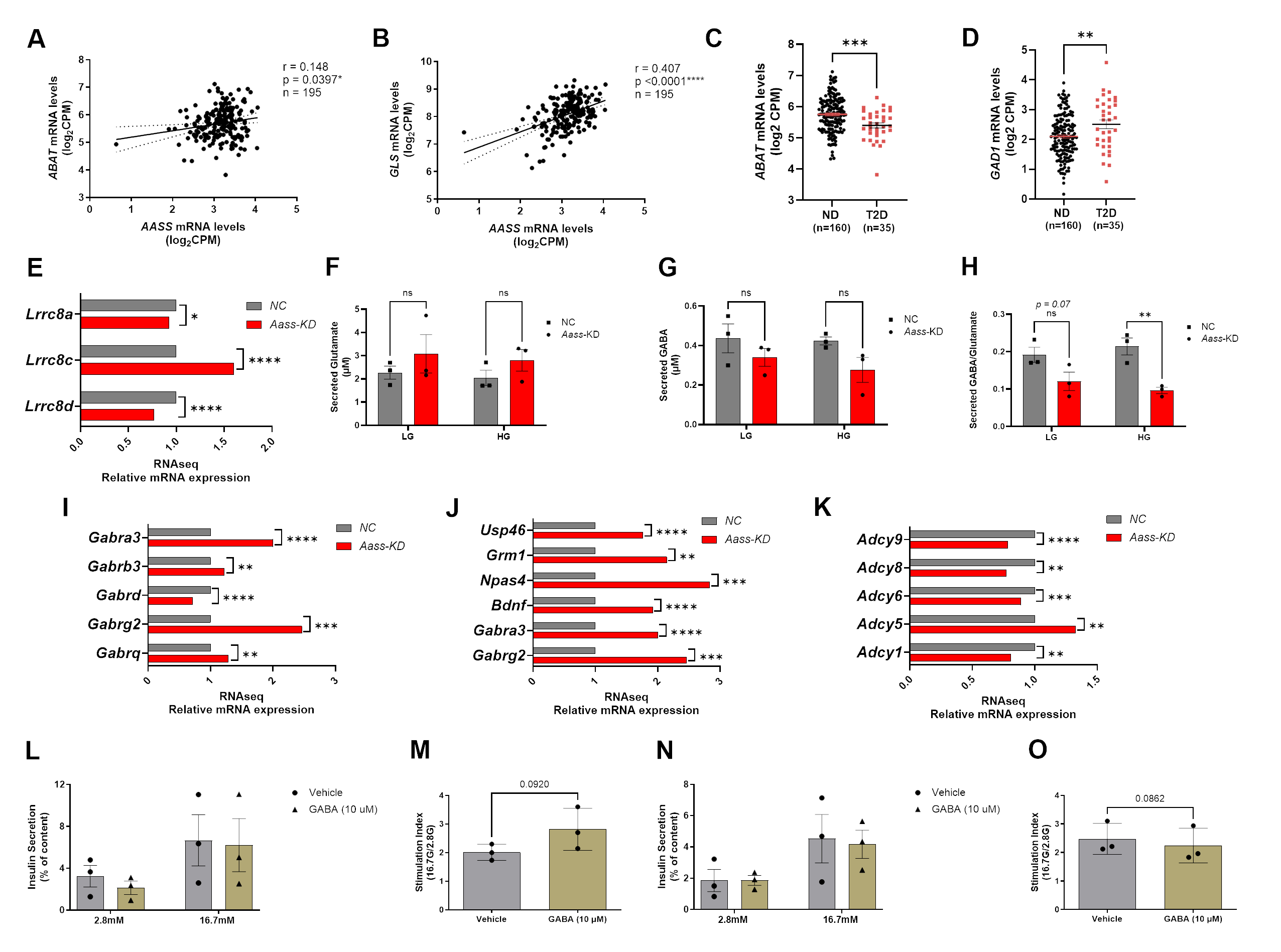

### Figure S4

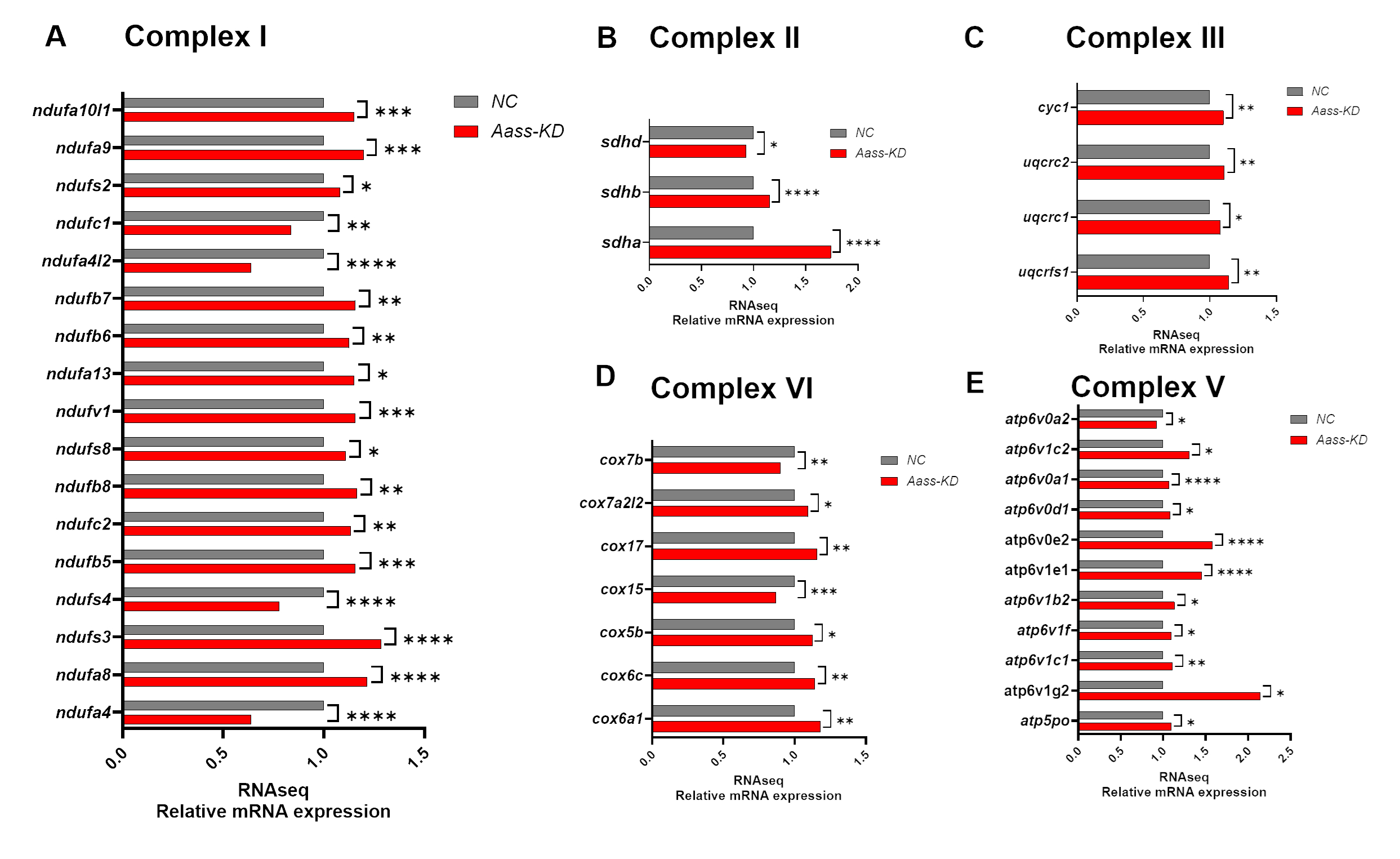
